# Mapping EGFR1 sorting domains in endosomes with a calibrated 3D expansion microscopy toolkit

**DOI:** 10.1101/2025.10.01.678490

**Authors:** Tayla Shakespeare, Rajpinder S. Seehra, Neftali Flores Rodriguez, Nkolika Atuanya, Thomas M.D. Sheard, Ralf Köhler, Daniel Bose, Philip Woodman, Barbara Ciani, Izzy Jayasinghe

## Abstract

Endosomes are nanoscale intracellular compartments that sort and recycle cell surface receptors such as epidermal growth factor receptor 1 (EGFR1). Nanometre-scale interactions and co-clustering of signalling proteins, cargo, and the membrane are critical to this process. Direct visualisation of these interactions has been hindered by the limited 3D resolution achievable with conventional and super-resolution microscopies. Here, we present the adaptation of expansion microscopy (ExM) to visualise, and quantify nanoclusters of endosomal proteins of human retinal pigment epithelial (RPE-1) cells. A 3D distortion analysis was developed leveraging the Farneback optical flow principle for detecting anisotropies in the hydrogel expansion. Analysis of pre- and post-ExM image volumes for 3D anisotropies revealed under-expansion of cytoplasmic regions within ExM hydrogels, often leading to over-estimation of size and distance measurements of small compartments such as endosomes. A self-assembling protein nanocage that reports the true local and nanoscale expansion factor was genetically introduced into the cells to calibrate ExM images of cytoplasmic regions containing endosomes. To stimulate and visualise the internalisation and sorting of EGFR1 in mammalian cells, a pulse-chase protocol was carried out with fluorescently-tagged EGF. The cells were subsequently fixed at 15- and 30- minute time points and subjected to 10-fold ExM and multiplexed 3D Airyscan microscopy to map cargo and EGFR1 vs other endosomal proteins. A volume tracing pipeline was developed to visualise the changes in the labelled EGF and EGFR1 densities at the limiting membrane of the endosomes. With multiplexed 3D ExM image volumes, we observed the enrichment of both EGF and EGFR1 in the endosomal interior and the accumulation of endosomal protein Rab5a near the limiting membrane during this early maturation of the endosomes. Taken together, the multiplexed 3D ExM toolkit offers a quantitative framework for visualising and measuring the intrinsic biology of small sub-cellular organelles like endosomes at true molecular-scale resolution.

## Introduction

The spatial and temporal organisation of membrane proteins within the endosomal system is critical for regulating cellular si0gnalling, receptor trafficking, and membrane homeostasis. A prime example is Epidermal Growth Factor Receptor 1 (EGFR1), which, upon ligand binding at the plasma membrane, is rapidly internalised and trafficked through the endolysosomal system [1]. They are sorted via small endosomal compartments and recycled back to the plasmalemma or trafficked to the lysosomes for degradation. This precise sorting process represents a critical decision point where cells must balance between terminating persistent signals and maintaining cellular responsiveness by regulating the surface receptor pool. Endocytosed EGFR1s often assemble into discrete nanoscale domains or nanoclusters [2, 3]. These domains are enriched for activated receptor and can recruit specific sets of effector proteins that modulate signalling duration, strength, and pathway specificity.

Multiple signalling and effector proteins orchestrate the dynamics and fate of EGFR1 nanoclusters. Rab GTPases, such as Rab5A and Rab7A, define the temporal identity of early and late endosomes [4]. Rab5 activates the phosphoinositol 3-phosphate (PI3P) kinase Vps34 to generate local pools of PI3P [5], and mediates the recruitment of downstream factors including early endosome antigen 1 (EEA1) via direct interaction and through the binding of the EEA1 FYVE domain to PI3P [6]. PI3P helps recruit the ESCRT-0 component hepatocyte growth factor-regulated tyrosine kinase substrate (Hrs) to nanodomains that drive EGFR1 sorting to intraluminal vesicles (ILVs) [7, 8] The FYVE-domain protein, endofin (also known as ZFYVE16), further enriches these nanodomains, coupling phosphoinositide recognition to SMAD signalling and ILV sorting [9, 10]. The interactions of endosomal proteins are spatially regulated, yet their spatial organisation and the nanoscale architectures of endosomes facilitating this process remain challenging to visualise with conventional microscopies.

Endosomes are typically 100–300 nm in diameter across their limiting membranes; many of the finer membrane topologies such as tubules and intraluminal vesicles are only tens of nanometres in scale [11, 12]. Diffraction-limited optical microscopy (with lateral and axial resolution limits of ∼250 nm and ∼600 nm, respectively) cannot resolve individual endosomes, nor can it distinguish protein nanoclusters that may be separated by <100 nm. Even conventional super-resolution techniques such as STORM, PALM, or SIM, whilst offering improved spatial resolution, remain inherently unsuitable for 3D volumetric imaging of densely packed cellular compartments. 3D implementations of STORM or PALM, such as biplane or astigmatism-based methods, could improve the axial resolution to ∼50–70 nm, allowing the visualisation of protein nano-clusters around small organelles such as clathrin-coated pits [13]. Alternatively, combining of 2D protein localisation with tomographic electron microscopy (superCLEM) provides a ground-truth for molecular localisation of protein clusters and cargo into sub-endosomal domains such as recycling tubules that may return the proteins to the plasmalemma [11]. Whilst sub-10-nm localisation techniques such as DNA-PAINT have emerged as powerful tools for multiplexed analysis of endosomal markers [14], they remain best suited for near-field imaging of flat and/or relatively sparsely labelled samples. These type of image data therefore remain principally limited to 2D.

Expansion microscopy (ExM) overcomes many of these limitations by physically inflating the sample prior to imaging, decoupling resolution from the limits posed by diffraction [15]. When combined with confocal microscopy, ExM enables isotropic resolution below ∼ 70 nm [16], even in optically-thick samples such as tissue sections and whole organisms [17-19]. With structured illumination or Airy-scanning imaging, effective resolutions of 15–30 nm can be routinely achieved [16, 20]. Critically, ExM enhances both lateral and axial resolution, making it ideally suited for resolving nanoclusters within volumetric compartments such as endosomes. Variations of ExM chemistry over the past decade have also allowed different degrees of expansion, differential targeting of biomolecule classes (e.g. proteins, lipids, or nucleic acids) [21-23], and/or different degrees of expansion isotropy [24, 25].

A key barrier to widespread adoption of ExM for quantitative molecular imaging lies in its spatial fidelity. Physical expansion can be heterogeneous, leading to distortions and variable expansion factors (EFs) across regions of interest. Errors in ExM can manifest either as distortions of the sample ultrastructure or under-expansion [26] that can limit the quantitative nature of ExM image data. Emerging solutions for this include registration of post-ExM images against an pre-ExM image [27, 28], imprinting the gels with fiduciary patterns [29], and statistical analysis of intrinsic ultrastructures such as microtubules and sarcomeres [20, 30]. However, the lack of a truly nanoscale reporter of the expansion of protein-rich cellular ultrastructure has remained a key barrier to its adoption for visualising subcellular compartments.

In this paper, we present a quantitative framework for 3D ExM imaging of small intracellular compartments with a focus on the nanoscale organization of EGFR1 sorting within the endosomal system. To verify the degree of expansion of sub-cellular structures, we have developed a 3D error detection analysis pipeline and adapted a genetically encoded self-assembling protein nanocage as an intrinsic calibrant to the local EF. By leveraging this quantitative 3D ExM imaging approach, we have mapped the distribution of activated EGFR1 and EGF ligands within the maturing endosome relative to key sorting effectors, including Rab5A and endofin at specific time points following endocytosis.

## Results

### 3D expansion microscopy for imaging endosomal proteins

We adopted ExM for visualising the 3D organisation of endosomal proteins within human Retinal Pigment Epithelial (RPE-1) cells. Proteins associating the endosomal membrane, immunolabelled *in situ*, were subjected to the expansion protocol which consisted of the principal steps of anchoring and gelation, digestion, and the subsequent expansion by hydration with deionised water (dH_2_O; schematically summarised in Figure 1A). In Airyscan images of unexpanded cells, endosomes labelled for early endosomal antigen-1 (EEA1) commonly appeared as bright, diffraction-limited puncta in perinuclear regions (Figure 1B). By comparison, similar cell samples expanded with 10-fold (10x) ExM and imaged with the resolution-enhancing Airyscan [31] featured noticeably more torus-shaped labelling patterns that reflected the endosomal surface localisation of EEA1 (Figure 1C). On close examination of images, we confirmed that the smaller endosomes (Ø <200 nm) were unresolved whilst a ring-like morphology was present only in larger (Ø >300 nm) endosomes in Airyscan images of unexpanded samples (magnified views shown in Figures D-i & E-i). In samples expanded with 4x ExM and imaged with Airyscan, the luminal spaces of both small and large endosomes were discernible. However, the boundary of EEA1 labelling on the endosomal surface appeared dense and continuous (Figures D-ii & E-ii). By comparison, in samples expanded 10-fold, the EEA1 labelling followed a more discernibly punctate morphology on the endosomal surface (Figures D-iii & E-iii). This punctate morphology resembled either individual markers or small clusters of well-resolved individual antibody markers observed before in other cell types by combining 10x ExM with Airyscan [20].

**Figure 1.**
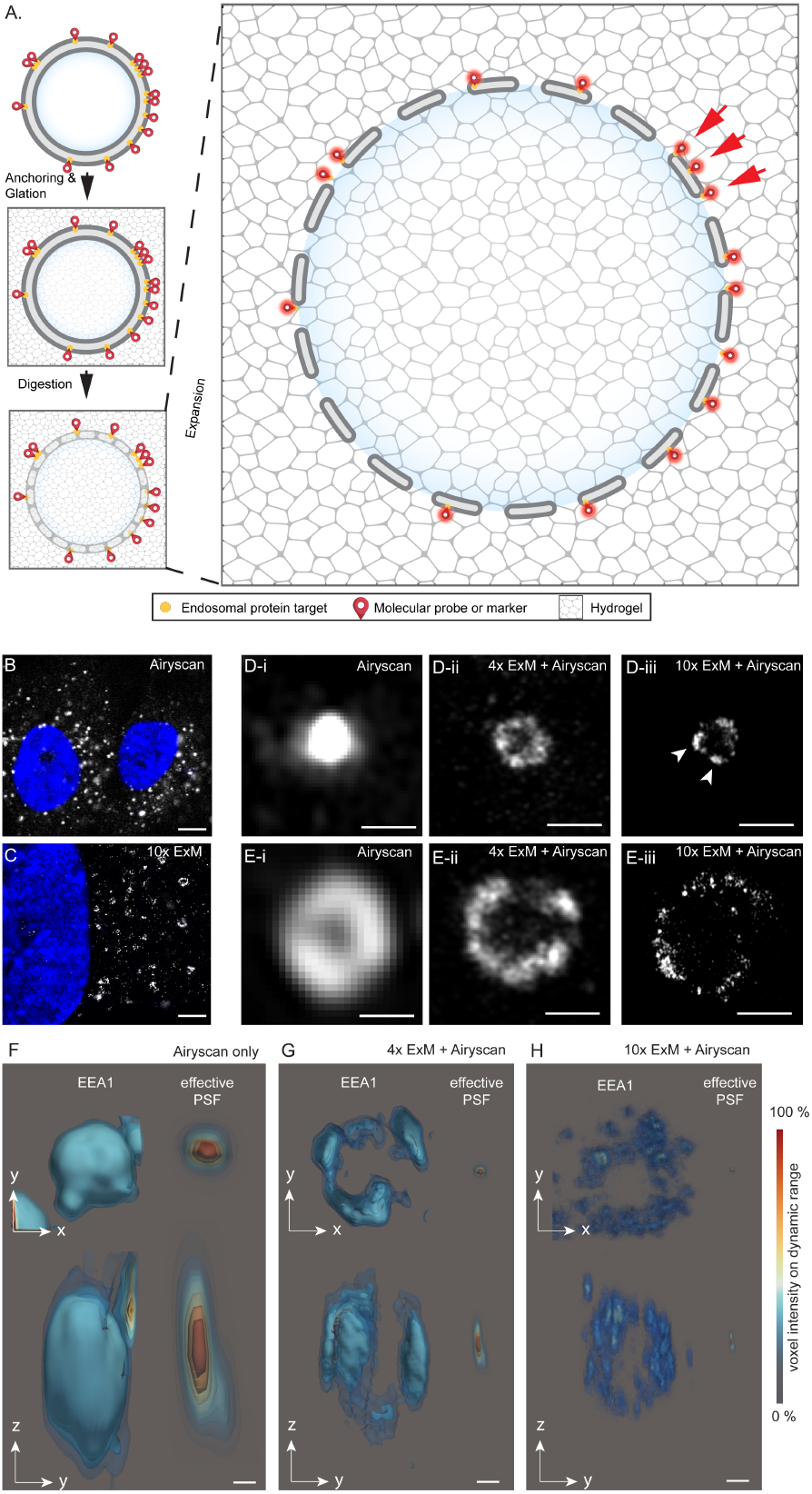
3D expansion microscopy of endosomes. **A.** Schematic diagram of ExM of endosomal proteins. In this method, protein targets in the cells are labelled and chemically anchored before the hydrogel is polymerised *in situ*. Following a mild enzymatic digestion, the molecular-scale structure and markers imprinted onto the hydrogel matrix is expanded by osmotic swelling to physically inflate the samples. Previously unresolvable structures are now resolved due to the physical separation of their markers (red arrows). **B**. Confocal image of un-expanded RPE-1 cells labelled for EEA1 (grey) and nuclear stain DAPI (blue). **C**. Airyscan image of an identical sample following 10-fold expansion (10x ExM) reveals the EEA1 endosomes as ring-like structures compared to diffraction-limited puncta. Magnified confocal images of **(D-i)** small and **(E-i)** large endosomal vesicles (Ø <100 nm and >300 nm respectively), labelled for EEA1 illustrate the poorly resolved nature of the spherical endosomal shapes. By comparison, 4x ExM Airyscan images show both **(D-ii)** small and **(E-ii)** large endosomes of equivalent sizes as torus-shaped morphologies. With the superior resolution achieved with 10x ExM combined with Airyscan nanoclusters (arrowheads) of EEA1 labelling were observed on the surfaces of both **(D-iii)** small and **(E-iii)** large endosomes. 3D isosurface contour rendered examples of endosomes from **(F)** unexpanded, **(G**) 4x ExM, and **(H)** 10x ExM RPE-1 cells labelled for EEA1 are shown in both x-y (upper) and z-y (lower) views. Shown on right in each panel, is the x-y and z-y view of the effective PSF (relative to the scale-corrected endosomes) as a result of the expansion. Colour scale reflects % of the voxel intensity across the full dynamic range of the structure. Scale bars: B: 5 μm, C: 2 μm D&E: 200 nm, F-H: 100 nm.

The resolution improvement achieved from ExM is the result of an effective downscaling of the point spread function (PSF). To estimate the PSF of the Airyscan protocol used in our experiments, we imaged polystyrene microspheres with Ø 100 nm, averaged between 10 copies. Examples of exemplar endosomes from RPE-1 cells are compared between unexpanded, 4x ExM, and 10x ExM (Figure 1 F-H). To scale, the effective PSF is shown in each example. This comparison reveals that the nanoclustering morphology on the endosomes, both on the top and lateral sides is only resolved as the PSF is scaled below the typical size of each nanocluster. This is principally achieved with 10x ExM, as the effective size (reflected by the full-width at half-maximum; FWHM) of the PSF approaches ∼17.5 nm laterally, and ∼65 nm axially. Features of the Airyscan volumes of 10x ExM samples was the hollow interior of endosomes and the intricate patterns of EEA1 nanoclustering visible on both bottom, lateral, and top surfaces (Supplementary Figure S1). Whilst the intensity and density of the lateral nanoclusters was higher due to the poorer axial resolution and the greater axial signal integration, the pattern of nanoclusters on the top surface was best resolved in glancing, in-plane optical sections of the bottom surfaces of these endosomes (Supplementary Figures S1 & S2).

### Tools for spatial error analysis in ExM of endosomes

A key limitation of ExM is that anisotropic gel expansion can lead to distortions of the ultrastructure and/or incorrect estimation of the EF in the region of interest. To observe distortions, ExM hydrogels of RPE-1 samples immunolabelled for EEA1 within bespoke, geometry-preserving microplates [26] were imaged both prior to, and after expansion. The square geometry of the hydrogels polymerised and expanded within each well of the microplate allowed us to record x-y coordinates of cells and structures of interest prior to expansion, and then to efficiently track and re-image the same structure post-expansion (See details in Supplementary Methods section 3.1). Figure 2A illustrates scaled and aligned 2D overlays of pre- and post-expansion Airyscan images (red and cyan respectively) of a cell subjected to a 4x ExM experiment. The overlaid field of arrowheads report the local distortion shift fields in the x-y planes. Whilst most endosomes aligned with high fidelity (e.g. Figure 2A-i), some endosomes in the same image appeared more significantly shifted (Figure 2A-ii; arrows indicate shift vectors). The aligned images were sub-sampled and the root-mean-square of the error (RMSE) was calculated from the resultant distortion shift vectors for each sampling scale. Figure 2B plots the RMSE across different length scales (where distance values reflect the post-ExM scale) ranging from 50 nm to 2.5 μm (Figure 2B) indicating distortion error of ∼ 1-5 % in the typical measurement range for 200-1000 nm.

**Figure 2.**
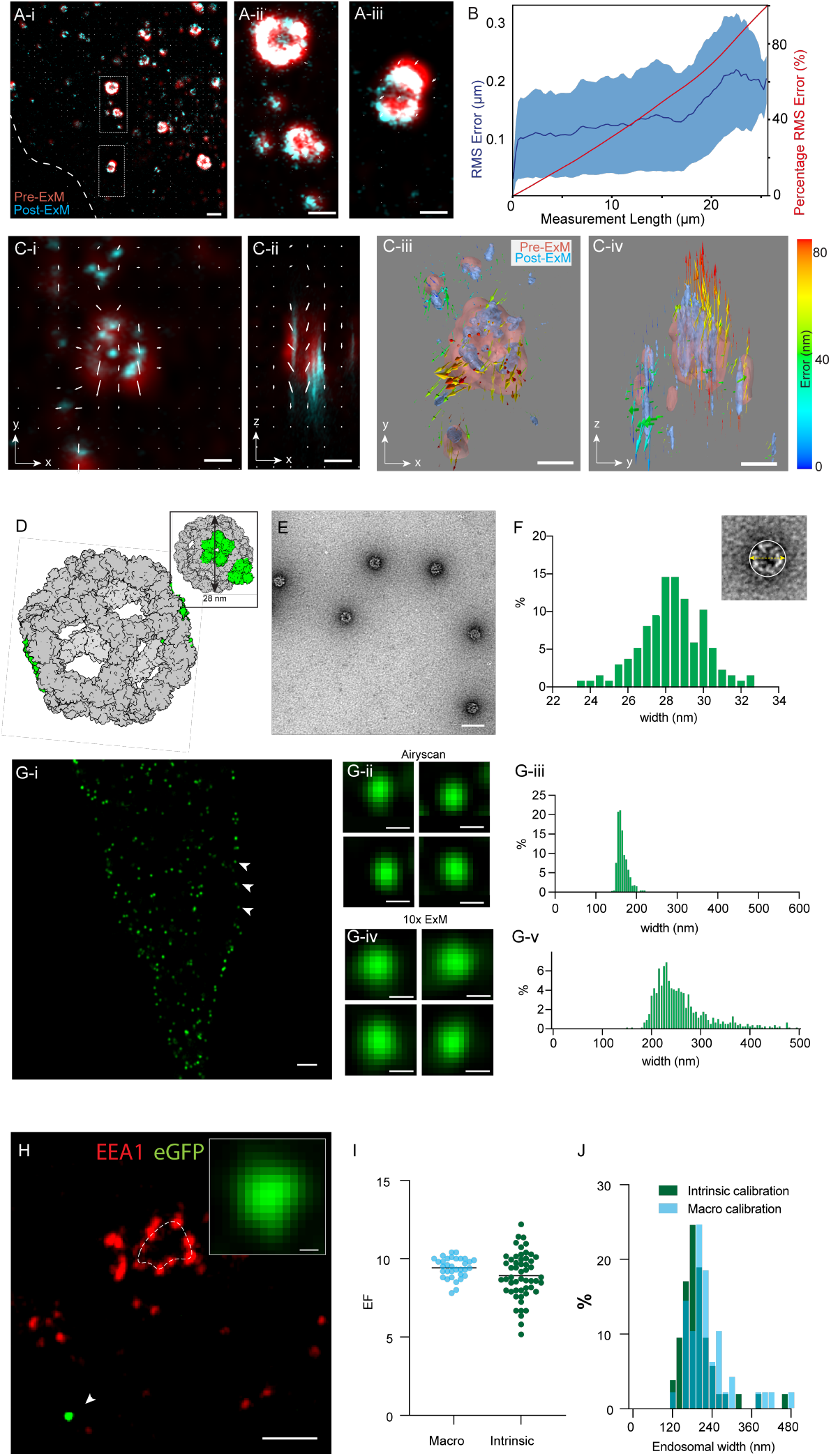
Calibrated expansion microscopy of endosomes. **A-i.** Image of EEA1 labelling in a RPE-1 cell shows pre- (red) and post-ExM (cyan) image is overlaid with the distortion vector field (white arrows indicating magnitude and directions of shift). Magnified views of endosomes indicate most endosomes showing negligible registration errors (**A-ii**) whilst a minority or regions indicate shifts ∼ 1 % in magnitude (**A-iii**). **B**. The plots of RMS error (mean curve across 6 datasets shown in dark blue; SD in light blue shading) and cumulative percentage error at different length scales of the image derived from this distortion analysis. **C-i & C-ii** are x-y and x-z views of a volumetric alignment of an endosome imaged before (red) and after (cyan) 10x ExM. The white arrows indicate the distortion vector components in each view. **C-iii & C-iv** illustrate isosurface rendering of the same volume from the same respective points of view. The colour scale (right) indicates the magnitude of each distortion vector. **D**. To report intrinsic expansion factor, a self-assembling decahedral 60-subunit nanocage (grey) with two sfGFP per subunit (green) with an overall estimated diameter of ∼ 28 nm were expressed in the cells. **E**. Negative-stain electron micrograph of purified nanocages indicating their self-assembly. **F**. A percentage histogram of the diameter of nanocages estimated with negative stain EM (illustrated as per inset). **G-i**. Fluorescence Airyscan micrograph of RPE-1 cell (boundary indicated by dashed line; DAPI stained nucleus in blue) expressing individual nanocages (green; arrowhead). Compared, are magnified views of exemplar nanocages image with Airyscan microscopy, pre-ExM (**G-ii**) and post-ExM (**G-iv**). Histograms illustrate width analysis of the nanocages in unexpanded Airyscan images (**G-iii**) and post-ExM Airyscan images (**G-v**). **H**. A 10x ExM image of a RPE-1 cell stained for EEA1 (red) observed adjacent to intrinsically expressed nanocage (green; magnified view in inset). **I**. Dot plot comparing estimates of expansion factor (EF) by macroscopic measurements (mean ± SD: 9.4 ± 0.6; n = 33 replicates), and intrinsic measurements based on mean nanocage widths (mean ± SD: 8.9 ± 1.4; n = 53 replicates). **J**. Overlaid histograms of estimated endosomal width from data calibrated for macroscopic EF (cyan) and intrinsic EF (green). Scale bars: A-i: 200 nm, A-ii & A-iii: 100 nm, C&D: 100 nm, E: 50 nm, G: 200 nm, H: 200 nm, K-inset: 10 nm

To further investigate the 3D nature of the distortions, arising particularly in experiments with higher EF, we developed a 3D distortion mapping approach. A block-matching approach for calculating a transformation matrix was used for rescaling and aligning the pre- and post-ExM image volumes of the same region of interest (See details in Supplementary Methods section 3.2; Supplementary Figure S9). Registered image volumes were re-sectioned into single 2D planes, both in x-y and x-z. An adaptation of Farneback’s optical flow technique allowed us to calculate the 2D distortion shift vectors for a given point in both x-y and x-z planes (see details in Supplementary Methods section 3.3). Figure 2C illustrates in-plane (x-y, Figure 2C-i) and an axial (z-x; Figure 2C-ii) views of the pre- and post-ExM volumes containing one endosome (red and cyan respectively), overlaid with the local distortion shift vectors (white arrows). The orthogonal shift components were used for calculating the true vectors, shown as arrows in Figure 2C-iii & iv. Radially oriented shift vectors (also see Supplementary Figure S3) were commonly observed at the endosome in spite of good registration of the overall cell. Localised alignment error reflected under-expansion of the small compartments like endosomes, post-ExM (shown with a blue isosurface in panels C-i to iv), compared to the pre-ExM features (red).

### Protein nanoscale reporter of intrinsic gel expansion

The sensitivity of this approach of distortion detection is ultimately limited by the resolution of the pre-ExM image. A truly nano-scale reporter of cellular expansion is required to verify cytosolic under-expansion. We adapted a homomultimeric 60-subunit decahedral protein nanocage developed previously by David Baker’s laboratory [32] (Figure 2D) as a calibrant of the local gel expansion. The cDNA for the I3-01_60eGFP monomer of the nanocage was cloned onto a mammalian vector before transfecting the RPE-1 cells which then expressed each subunit of the nanocage consisted of eGFP, self-assembled into a 60-meric nanocage with a Ø ≈ 28 nm (inset), observable throughout its cytoplasm. Negative stain transmission electron microscopy (TEM) was used to measure the widths of individual nanocages isolated and purified from similar cells (Supplementary figure S4). These measurements were comparable to nanocages purified and filtered from *E. coli* expression systems (Figure 2E, Supplementary Figure S4). The histogram in Figure 2F illustrate the distribution of the widths of purified nanocages (mean ± SD: 28.3 ± 1.6 nm; n = 138 nanocages; examples of TEM images at different stages of purification shown in Supplementary Figure S5). In Airyscan images of unexpanded RPE-1 cells, the eGFP fluorescence of the nanocages was observed as bright and diffraction-limited spots throughout the cell (arrowheads; Figure 2G-i). Close examination of the spots in these Airyscan images (magnified view of examples shown in Figure 2G-ii) and the subsequent width measurements (histogram in G-iii with a mean ± SD of 165.6 ± 12.8 (n=678 nanocages; 3 cells)) confirmed this. Analysis of the cytoplasmic nanocages in Airyscan images following 10x ExM revealed larger, supra-resolution spots (Figure G-iv & v), and a broader distribution of widths (mean ± SD: 263.7 ± 63.1 nm; n = 794 nanocages; 7 cells). The latter was consistent with local heterogeneity in the expansion of the nanocages throughout the cell volume. When the transfected cells were immunolabelled for EEA1, the nanocages were commonly observed in regions adjacent to endosomes, and within the same image planes (Figure 2H). To calibrate the local average EF we divided the mean width of the nanocages in post-ExM Airyscan images by the mean width of the nanocages estimated with TEM. A comparison of the intrinsic EF estimates based on the nanocage measurements against the macroscopic EF estimates (by measuring the dimensions of the hydrogel before and after 10x ExM expansion) across 33 samples is shown in the dot plot in Figure 2I. Whilst the intrinsic EF measurements displayed greater variability (SD of 1.4 compared to 0.6), it also showed ∼ 6% lower mean (p = 0.04; df =86; one-tailed Mann-Whiteney U-test).

Figure 2J illustrates the left-shift in a histogram of endosomal widths (a median drop of 20.3 nm; p <0.05, df = 100; Wilcoxon test) measured from 10x ExM Airyscan data once the intrinsic EF was used to calibrate the image instead of the macroscopic EF (see similar analysis for 2D area of endosomes in Supplementary Figure S6).

### Visualisation of EGF and EGFR1 sorting using ExM

To visualise endosomal sorting of EGFR1, we performed a series of pulse-chase experiments employing Alexafluor^488^-conjugated streptavidin bound to biotinylated EGF ligand. The cells were fixed at 15 min and 30 min time points following the EGF pulse, allowing the internalised EGF/EGFR1 complexes to be sorted via early endosomes and their maturation into multivesicular bodies (Figure 3A). Following fixation, EGFR1 was additionally immunolabelled, allowing us to simultaneously visualise the spatial distribution of the receptor and the ligand (Figure 3B). Unexpanded Airyscan images showed bright, diffraction-limited spots of EGFR1 labelling (magenta) throughout the cytoplasm of the cells. Closer examination of magnified images revealed co-localisation of discrete EGF puncta with the EGFR1 outline of larger endosomes. ExM Airyscan images of endosomes from cells fixed at 15 min revealed discrete nanoscale domains of EGFR1 clustering along the boundary of the endosome (indicated with dashed line; Figure 3E-i). Both EGFR1 and EGF were often localised to the vesicular interior in ExM Airyscan images of endosomes, particularly at 30 min (Figure 3F-i). 3D datasets of the 15 min time point were isosurface rendered and visualised in orthogonal views (Figure 3E-ii vs 3E-iii) revealing the overwhelming localisation of EGF and EGFR1 to the endosomal surface. By comparison, ExM Airyscan volumes from 30 min clearly indicates a greater subset of EGF/EGFR1 nanoclusters localised in the centre of the vesicle (example shown in orthogonal views in F-ii vs F-iii). To analyse the spatial relationships between the ligand and receptor labels, calibrated ExM Airyscan datasets were subjected to a segmentation using Huygens software, allowing us to discretise and localise individual puncta in both EGF and EGFR1 channels (illustrated schematically in Figure 3G). The violin plot comparing the measured widths of the segmented puncta (Figure 3H) led to two key observations from this analysis. Firstly, the EGFR1 puncta were consistently wider than the EGF puncta at both time points (by 84% at 15 min, and by 34% at 30 min). Secondly, the EGF puncta at the 30 min time point were ∼ 38% wider than that at 15 min (Mean ± SD of 44 ± 25 nm vs 32 ± 22 nm), particularly observed in the endosomal interior. Analysis of the nearest neighbour distance of each EGF-to-EGFR1 puncta revealed an ∼ 60% increase in the ligand-receptor marker distances from 42 ± 26 nm to 67 ± 30 nm between 15 and 30 min time points (Figure 3I). In contrast, the ratios between the total volume (Figure 3J) and the total number of puncta (Figure 3K) were consistently lower and less variable at the 30 min point, indicating a progressive enrichment of ligand-bound EGFR1 within the endosome as well as possible recycling out of the endosome.

**Figure 3.**
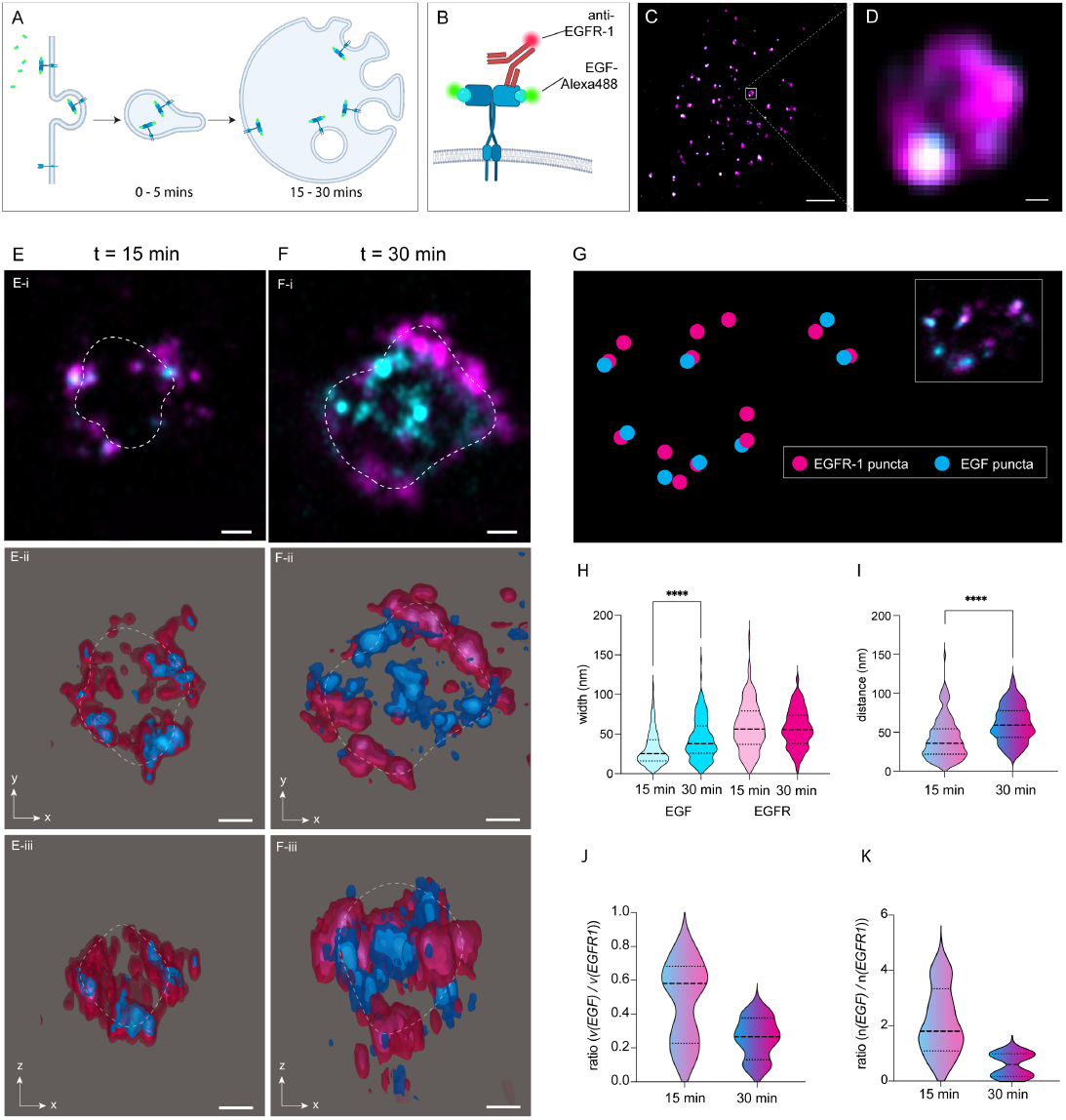
Nanoscale measurement of EGFR1 sorting during endosomal maturation. **A.** Schematic illustration of EGF stimulation leading to endocytosis of the EGF ligand-EGFR1 receptor complex into early, small endosomes and the subsequent sorting of EGFR1 during the maturation and expanding endosome over the timescales of 15-30 min (t). **B**. Illustration of the fluorescent pulse-chase dual imaging of the Alexa 488-EGF and post-hoc immunolabelled EGFR1 receptor allowing independent localisation of EGF and EGFR1. **C**. Airyscan overview image of a cell with dual EGF-EGFR1 labelling (cyan and magenta respectively). **D**. Magnified view of an endosome in panel C, exemplary of the toroid morphology of EGFR1 staining (magenta) and confined and co-localised pucta of EGF (cyan). 10x ExM images of EGF and EGFR1 at exemplar endosomes at pulse chase time points of (**E-i**) 15 min and (**F-i**) 30 min, noting the additional densities of EGF and EGFR1 localisation in the middle of the vesicle at the latter. The dashed white line indicates the approximated boundary of the endosomal labelling. Orthogonal views of isosurface visualisation of EGF (blue) and EGFR1 (magenta) of the same endosomes from 15 min (**E-ii & iii**) and 30 min (**F-ii & iii**) time points. **G**. To analyse the relationship between puncta of EGF and EGFR1, the punctate densities were segmented and the nearest neighbour distance from each EGF punctum to the nearest EGFR1 punctum analysed (corresponding original image shown in inset). **H**. Violin plot comparing the width of the EGF (cyan) and EGFR1 (magenta) puncta at 15 min (light coloured) and 30 min (dark coloured) time points. Comparisons: **** from Mann-Whitney test. p < 0.0001, df= 696 puncta. **I**. Violin plot of he measured nearest neighbour distance from EGF puncta to nearest EGFR1 **** p <0.0001, df = 213 puncta. Also shown, are violin plots of (**J**.) the ratio between the total volume of EGF puncta and the total volume of EGFR1 puncta and (**K**.) the ratio between the number of EGF and EGFR1 puncta in each endosome at 15 mins (n=9 endosomes, 9 cells) and 30 mins (n=9 endosomes, 8 cells). Scale bars: 50 nm.

### Multiplexed, 3D ExM of ligand, receptor, and sorting proteins

Rab5a is recruited to the endosomal surface and organised into nanodomains by the Rab5 guanine nucleotide exchange factor (GEF) complex of Rabex-5 and Rabaptin-5 [33-35]. Similarly, endofin is recruited to nanodomains of early endosomes that also contain ESCRT-0 and the ESCRT accessory factor HD-PTP, which collectively drive the sequestration of EGFR1 prior to its incorporation into ILVs [7, 10, 36, 37]. To examine the nanoscale relationship of Rab5a and endofin with EGF and EGFR1, we carried out a series of 3-colour 10x ExM Airyscan imaging experiments. Alongside the fluorescently-tagged EGF, we performed a four-way analysis between EGFR1, Rab5a, and endofin. Figure 4A and 4B illustrate overlays of EGFR1, EGF, and endofin in exemplar endosomes imaged at 15 min and 30 min time points. A similar comparison between EGF, endofin, and Rab5a (Figure 4C & D) demonstrated that the two latter molecules remain at the endosome surface.

**Figure 4.**
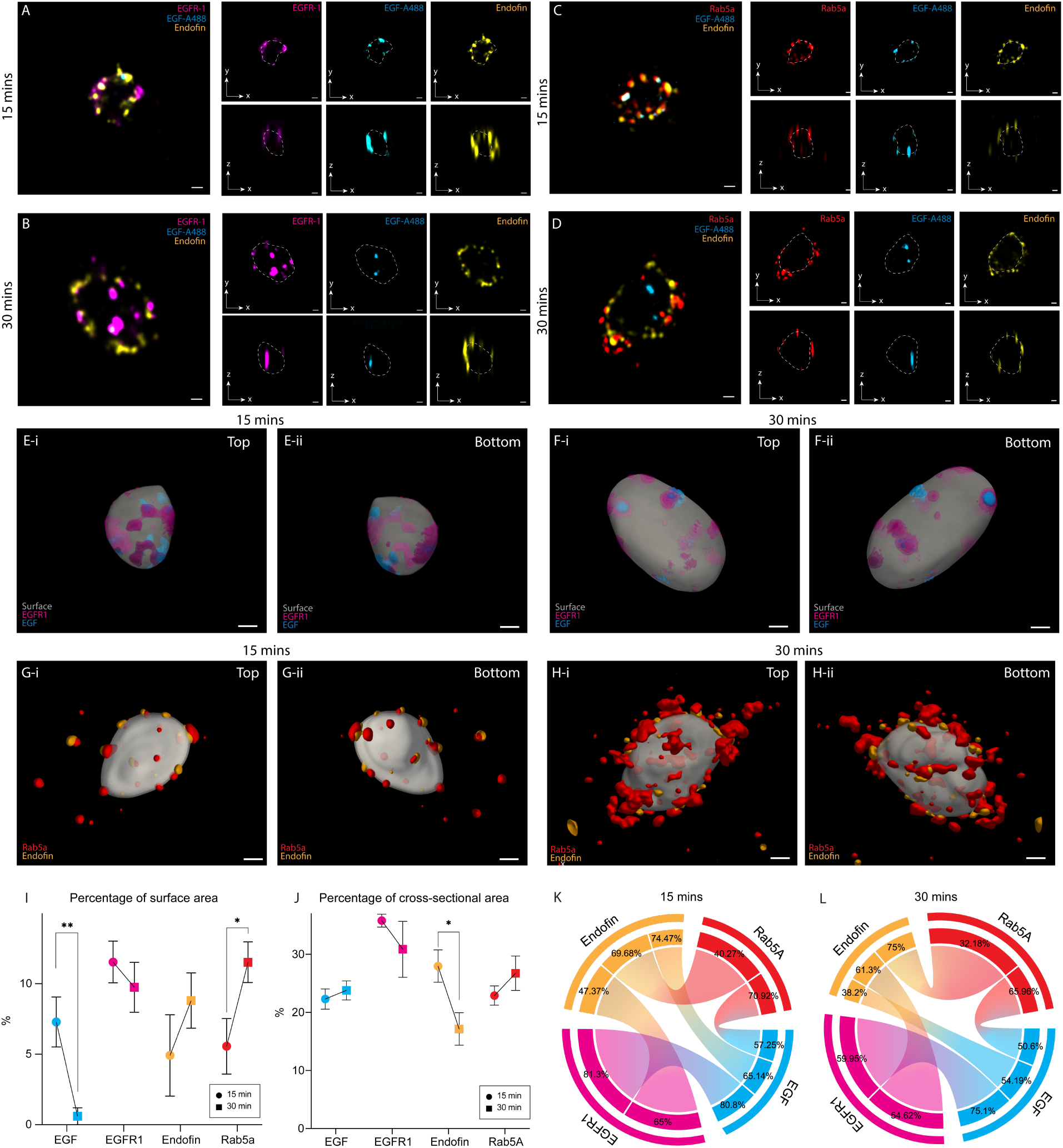
Visualisation of the 3D organisation of EGFR1 sorting proteins. Shown in the first comparison, are three-colour 10x ExM Airyscan image volumes overlaying EGF (cyan) EGFR1 (magenta), and endofin (yellow) labels in exemplar endosomes from (**A**.) 15 min and (**B**.)30 min time points following the Pulse-chase protocol. In the second comparison, equivalent image volumes overlaying EGF (cyan) Rab5a (red), and endofin (yellow) labels are shown at from (**C**.)15 min and (**D**.)30 min time points. In each example, individual channels of the volumes are shown in both in-focus x-y (upper row) and x-z (lower row) view. Top and bottom view of isosurfaces of volume-rendered examples of endosomes from (**E-i. & ii**.) 15 min and (**F-i. & ii**.) 30 min are shown. Translucent grey depicts the surface of the endosome traced based on endofin labelling whilst EGF (cyan blue) and EGFR-1 nanodomains are projected onto the surface. Opaque, and lighter coloured regions indicate the nanodomains in the reverse side of the endosome. Volume-traced endosomes from (**G-i. & ii**.) 15 min and (**H-i. & ii**.) 30 min timepoints isosurface rendered to show traced surface (translucent grey), endofin (yellow), and Rab5a (red), particularly its heterogeneous macro-clustering pattern at 30 min. (**I**.) Line plots of the mean percentage of endosomal surface area occupied by projected EGF and EGFR1 nanodomains (as shown in panels 4E and 4F) as well as endofin and Rab5a nanodomains at 15 min to 30 min time points. Error bars indicate SD. ** p<0.0001, df = 7 data sets; * p<0.001, df = 8 data sets. (**J**.) Line plots of the mean percentage of pixels of representing the cross-sectional area of the endosome occupied by each target of interest at 15 min to 30 min. Error bars indicate SD. * p <0.05, df = 8. Chord diagrams summarising the % of colocalisation of a protein of interest with the reference protein at (**K**.) 15 min and (**L**.) 30 min time points. For example, the % of Endofin markers overlapping with Rab5a at 15 min is 69.68%. The relative length of the arcs of each protein target of interest has been scaled in proportion to the 2D area occupied by the protein (summarised in panel I) at the corresponding time point. Scale bars: 100 nm.

To visualise the nanodomains occupied by these molecules, we developed a new method for tracing the endosomal volume that produced a model of its limiting membrane (See supplementary methods section 3.5). This method used the punctate labelling densities that line the compartment’s limiting membrane to construct an iterative series of Delaunay triangulation transformations which were then averaged to achieve a smoothed boundary of the endosome (Supplementary Figures S12 and S13). In Figure 4E and 4F, the adjacent EGFR1 (magenta) and EGF (cyan blue) labelling (within ± 40 nm of the boundary) were projected onto the traced endosomal surface (grey) to visualise the surface nanodomains they occupy (See Supplementary movie 1). A similar rendering of the endosomal surface show the 3D labelling densities of both Rab5a (red) and endofin (yellow) are initially recruited to distinct nanoclusters on the endosome surface (at 15 min; Figure 4G). Whilst the endofin puncta appeared to be relatively sparse at the endosomal surface at 30 min, larger densities (macro-clusters) of Rab5a were observable adjacent to the endosomal surface (Figure 4H; Supplementary movie 2).

Plots of the percentage of the endosomal surface area occupied by the projected EGF indicate a ∼ 90% reduction in density between 15 and 30 min (p<0.01; df = 7 endosomes), particularly as the size and overall surface area of the endosome grows. A doubling of the percentage of endosomal surface area adjacent to the projected Rab5a is also seen, reflecting the enrichment of Rab5a into the macro-clusters observed above.The percentages of the endosomal cross-sectional area occupied by EGF, EGFR, Rab5a, and endofin at the two time points are plotted in Figure 4J. An ∼ 39% percent reduction in the density of endofin and an ∼ 17% increase in the density of Rab5a were observed. In the pairwise Manders co-localisation analyses for 15 and 30 mins summarised in Figure 4K and 4L respectively (see measured values in Supplementary Table S1), a 21.2% reduction of the EGF co-localising with EGFR is consistent with the reduction in the EGF:EGFR1 ratio of puncta in Figure 3. The reduction in endofin density observed at both the limiting membrane and the endosome cross-section between 15 and 30 min (Figure 4I & 4J) coincides with a 9% reduction of endofin labelling co-localising with each of EGF and EGFR. In spite of the increase in Rab5a density and its accumulation near the limiting membrane between these two time windows, its co-localisation with EGF and endofin is mostly unaltered.

## Discussion

### ExM as an endosome imaging method

We report the adaptation of variants of widely used 4x and 10x ExM methods for imaging endosomes. The data presented here of 10x ExM combined with resolution-enhancing Airy-scanning microscopy achieve a level of spatial detail of the endosome that surpasses previous observations from localisation modalities alone. In addition to the capacity to visualise both top and bottom surfaces of the endosomal vesicles, this has allowed us to localise, measure, and quantify the nanoclusters of EGF/EGFR1 complexes and signalling proteins contributing to the sorting process (Figures 3H-3K, 4K & 4H). It is also the first 3D optical data that allow measurements of sequestered and/or enriched nanoclusters of these proteins within a small compartment. The much improved 3D resolution with this approach has been pivotal for developing a new protocol for endosomal volume reconstruction and visualisation of EGFR1 nanodomains at the limiting membrane (e.g. Figure 4-E&F).

The resolution improvement achieved with these methods is still limited by the linkage error between the markers (antibody complexes and streptavidin-linked EGF ligands). Whilst post-expansion labelling [38] has emerged as a solution for minimising this error, we have found that the gains promised by this method (for example, the reduction in autofluorescence and improved antigenicity) are (i) modest for structurally simple samples like RPE-1 cell monolayers, and (ii) adds complexity to post-labelling washing steps in order to avoid non-specific background. More robust versions of ExM methods designed to reduce spatial error (e.g. Magnify or U-ExM, [19, 25]) may offer more consistent matching between intrinsic and macro-scale EFs. However, we note that the 10-fold ExM data presented in Figure 2C featuring radial distortions around endosomes were acquired from an ExM recipe derived from a method with improved anchoring and gelation efficiency [39]. The application of 3D error detection analysis is hence always advisable, in spite of the improved gel chemistries that are increasingly available to the cell biology community.

### Visualisation of EGF sorting protein interactions

Between 15 and 30 minutes, we observed a 10% reduction in the surface EGFR1 density, along with a 90% reduction in the density of EGF (Figure 4I). We also observed (i) an increase in EGF-EGFR1 puncta nearest neighbour distances (Figure 3I), and (ii) a modest (20%) decrease in the Manders co-localisation measurements between EGF and EGFR (Figure 4K). The latter two observations may reflect two possible features of the EGFR1 sorting process. The selective sequestration of ligand-bound EGFR1 receptors would enrich them within the interior [40], while it is also possible that some EGFR lacking Alexa-conjugated EGF may have acquired unlabelled EGF during the chase period. Alternatively, this could reflect either a true dissociation and/or degradation of the EGF/EGFR1 complex by the 30 min time point; however, the continued enrichment of Rab5a observed at these endosomes at the 30 min time point does not support this explanation. Endofin, a key component of the EGFR1 sorting nanodomains at the limiting membrane maintains a high co-localisation (∼ 47 – 61%) with EGFR1 and remains at the membrane at 30 min (Figure 4K). In contrast, the co-localisation of endofin with the fluorescently-conjugated EGF reduces substantially (from 74% to 38%) The 38% reduction in the % of the endosome’s cross-sectional area occupied by endofin is explained in part by the increase in the endosomal size may also reflect removal of endofin from the endosome as the sorting of EGFR1-EGF complexes to ILVs progresses. Rab5a continued to accumulate at the endosomal limiting membrane between the 15 and 30 min time points, forming macro-clusters at distinct regions of the endosomal surface, visually resembling the lipid rich sub-domains observed previously in HeLa cells with 2D dSTORM of Rab5c [11]. Whilst we did not measure a statistically significant change in the Rab5a puncta size, the observation of macro-clusters in ExM volume data and the doubling of its coverage over the approximated limited membrane support previous observations of continued Rab5 enrichment on the surface nanodomains [41-43]. We anticipate the macro-clusters of Rab5a to be re-mobilised as the endosome continues to mature.

### New tools for quantitative ExM

#### A non-destructive and nanoscale reporter of gel expansion

Ten years on from the introduction of ExM, the need for improving the reliability and standardisation of the post-ExM images, as well as their accurate interpretation, continues to be an intense area of focus. Existing methods of distortion detection or spatial analysis remain fundamentally limited by the spatial resolution of the reference (pre-ExM) images, limiting the method’s utility as a quantitative tool for imaging nanoscale structures. DNA origami [24] has been used for this purpose previously in cell-free samples. The adaptation of DNA calibrants for cell imaging however would require anchoring chemistry compatible with both nucleic acids and proteins. A fundamentally protein-based calibrant such as this nanocage is a more straightforward and authentic reporter of protein-retaining gel expansion. Analysis of intrinsic cellular structures such as muscle z-discs, microtubules, and nuclear pore complexes [20, 24, 44] is a popular method to quantify the intrinsic expansion isotropy or EF. A calibrant such as the decahedral nanocage adopted here also provides a known ground-truth of its dimensions, independent of cell type or physiology. As a well-characterised and self-assembling structure compatible with both mammalian and bacterial expression systems [32, 45], this nanocage would be compatible with a wide range of ExM experiments. Once the cells were transfected, it required no additional modifications to the labelling, gelation, or expansion steps of the ExM protocol. Unlike recent methods of gel anisotropy detection that require physical alteration of the sample either through photobleaching [46] or feature imprinting [29], a genetically-encoded nanocage provides an entirely non-destructive approach to estimating the EF intrinsic to the structures and region of interest.

The chosen nanocage is best suited for 10-fold ExM recipes when combined with Airyscan. It was not suitable for 4x ExM samples unless it could be combined with a super-resolution image acquisition modality such as STORM [47]. We achieved the highest efficiencies in delivering the nanocages to the cell interior through cDNA transfection rather than pre-mixing of purified nanocages with the gelation/monomer solution. A sparse, cytoplasmic distribution of the nanocages achieved this way was naturally the result of this expression approach. Due to the lack of compartment or cytoskeletal tether within the cell, some nanocages may be lost during the cell fixation or subsequent processing steps. The volumetric expansion of the cell during ExM (and the effective narrowing of the optical sectioning) also leads to a perception of sparsity in each Airyscan image plane. A greater density of nanocages could have provided a higher-resolution map of the gel isotropy. However, the natural sparsity observed in our samples was best suited for preventing aggregation and for avoiding cytotoxicity to the cell.

#### 3D distortion mapping and 3D reconstruction tools

Distortion mapping through pre- and post-ExM imaging has been the primary method of benchmarking expansion isotropy since the inception of the ExM concept [15]. Distortions resulting from expansion anisotropy are always three-dimensional. The 2D analyses available to-date only capture the distortion components in the x-y planes. In our method, the integration of the components in the orthogonal planes was critical for our discovery of the radial nature of the misregistrations local to some endosomes. For all users adopting ExM, irrespective of single-plane or volume imaging, this approach offers a more robust benchmarking tool over the 2D methods.

One of the remaining limitations of ExM is the limited repertoire of reliable membrane labels specific for small intracellular compartments that can survive gel expansion. In the endosomal volume reconstruction, endofin was used as an antigen marker of the limiting membrane. Similar to many membrane marker antigens, endofin’s punctate morphology only lends a discontinuous impression of the membrane. Our volume reconstruction approach of iteratively detecting and fitting puncta, and projecting them onto a triangulated surface, has helped us transform a relatively sparse, labelling pattern into a continuous volumetric representation of the vesicle. This method offers a scalable pipeline that can be translated and applied to ExM volumes of numerous other types of organelles or cells. In connection to endosomes however, this method carries two principal limitations. As an inherently smoothing-based approach, this algorithm is unsuitable for reconstructing finer topologies such as membrane tubules or invaginations unless they are discernible with a specific marker. Without specific markers, we were also unable to reconstruct intraluminal vesicles (ILVs) which carry out the sequestration of the EGF/EGFR1 complexes.

## Materialsand Methods

### Cell culture

Human retinal pigment epithelia RPE-1 cells were cultured in DMEM/F-12 Ham liquid with L-glutamine and sodium bicarbonate (Gibco D8062, Merck, UK) and 10% fetal bovine serum (FBS; v/v; Cat no. 17563595 Fisher Scientific Ltd, MA), supplemented with 1% Penicillin-Streptomycin (v/v; containing 5,000 units/mL of penicillin 5,000 μg/mL of streptomycin; Thermo Fisher, MA). For transfections in RPE-1 cells, Penicillin-Streptomycin was omitted from the cell culture medium for >1 passage. HeLa cells were culture in DMEM supplemented with 10% fetal bovine serum and supplemented with 1% Penicillin-Streptomycin. Cells were incubated at 37° C and 5% CO2 and seeded once they reached ∼ 70-80 % confluency, typically 2 days after passage. After removing the excess, cells were washed with PBS buffer and detached from flasks with 0.05 % Trypsin-EDTA (Cat no. 25300054, Fisher Scientific), and Trypsin was neutralised with complete fresh medium.

Cells were seeded onto No 1.5 glass coverslips (Epredia X1000 Cover glasses 22 x 22mm # 1.5 (0.16 – 0.19 mm), Cat no. 15805214, Fisher Scientific) in 6-well plates at a density of 50,000 cells /ml (2 ml of the passaged mixture). Seeded cells were incubated in cell culture medium at 37°C and 5 % CO2 for ∼ 2 days, until they reached ∼ 70-80% confluency. Intrinsic calibration of X10 ExM was performed in RPE-1 cells seeded at a density of 20-25,000 cells/ml (1 ml of the passaged mixture per well) and incubated for ∼ 2-3 days before transfection at ∼ 20-40% confluency. Cell culture medium was replaced with fresh complete medium (2ml per well) >4 hours before transfection or pulse-chase experiments.

Following transfection or pulse-chase experiments, coverslips were transferred to 2% PFA (v/v; 4% PFA solution in PBS was mixed 1:1 with 1% PBS to make up to the final concentration of 2%; Fisher Scientific Ltd) for 10 minutes at room temperature. Cells were washed 3x in 1% PBS to wash off excess PFA before quenching with ∼ 3-4 drops of 1M glycine in Tris buffer or 0.5% BSA (w/v, Gibco, Thermo Fisher Scientific) in a cell storage solution consisting of 1% PBS (tablets dissolved in dH2O; Fisher Scientific).

### EGF pulse-chase

RPE-1 cells were cultured on coverslips to ∼ 70 % confluency for ∼ 2 days as discussed above. Cells were serum starved for ∼4 hours in serum-free DMEM at 37°C 8% CO2. Cells were then incubated with 50 ng/µl EGF conjugated by a streptavidin linker to Alexa-488 (final [EGF] = 5 ng / µl) for 5 minutes at 37 °C 5% CO2 and coverslips were immediately transferred to serum free medium containing 50 ng / µl unlabelled EGF. Cells were incubated for 15 or 30 minutes before fixation and immunostaining.

### Expression of decahedral nanocages in cultured HeLa and RPE-1 cells

The mammalian expression vector encoding a single monomer subunit of the self-assembling 60mer nanocage (Table 1) developed by David Baker’s laboratory [32] was transfected for expression RPE-1 cells. Cells were cultured and seeded onto coverslips and maintained until they reached ∼20-40% confluency. They were were transfected with plasmid DNA using lipofectamine per manufacturer’s instructions, and optimised for cell viability and nanocage expression levels.

**Table 1.**
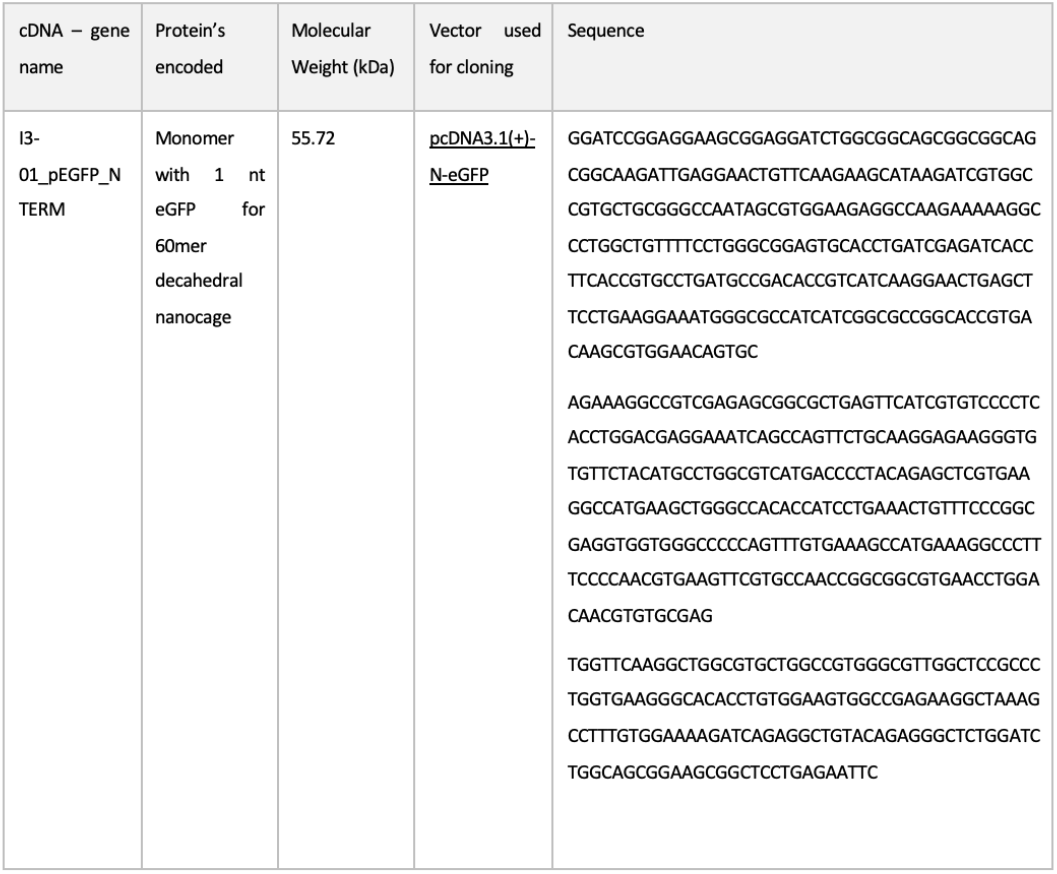
Details for cDNAs used to express nanocages in mammalian cells. The name, molecular weight and sequence is provided for mammalian expression vector for nanocage expression.

Cells were transfected with 0.5 µl plasmid DNA and 0.5 - 1.5 µl lipofectamine 2000 (Cat. 11668019 Thermo Fisher) per 10 cm2 well containing 2 ml cell culture medium. For each well, 0.5 µg plasmid DNA was gently mixed with opti-MEM medium (Cat. 31985062, Thermo Fisher) to a final volume of 125 µl. 0.5 µl or 1.5 µl Lipofectamine 2000 reagent was mixed gently with opti-MEM medium to a final volume of 125 µl. DNA and lipofectamine dilutions were incubated for >5 minutes at room temperature before mixing them together by gentle pipetting, and the resulting transfection mixture was incubated for >15 minutes at room temperature. 250 µl of the transfection mixture was added dropwise to plated cells in 2ml cell culture medium in a single well of a 6-well plate as described above. After 20-24 hours of transfection, cells were fixed and either stained with DAPI (diluted to 1 µg/ml in 1x PBS) for immunofluorescence imaging or expanded in ExM.

The DNA, PLUS reagent, and lipofectamine LTX or lipofectamine 2000 concentrations were adjusted in parallel depending on cell confluency. All transfection conditions gave similar results that varied between cells. Higher concentrations of transfection reagents improved expression but resulted in aggregation of nanocages in some cells, and lower concentrations resulted in even expression in some cells, but expression was limited in some cells. Cells were chosen for analysis based on the even expression of nanocages in the cell.

### Purification and characterisation of nanocages from mammalian cell lysate

HeLa cells expressing nanocages were washed with ice cold PBS (1x), and 1 ml of ice-cold lysis buffer (1 x PBS + 10 mM EDTA + 1 x protease inhibitor cocktail (Cat. S8830, Merck Life Sciences, Darmstadt, Germany) + 1 % Triton-X (Merck)) was added to cells plated on each coverslip. Lysed cells were scraped using a pipette tip and moved into centrifuge tubes. Lysates were kept on ice for 30 - 45 minutes with intermittent mixing by inverting. Lysates were span down at 13,300 rpm (17,000 x g) for 20 minutes at 4 °C. Supernatant was removed and concentrated through a 1MDa spin concentrator (Vivaspin 500, Merck) and stored at 4 °C until imaging with electron microscopy.

### Immunostaining

Following fixation, cells were stored in PBS containing 0.5% bovine serum albumin (w/v) or PBS with ∼ 3-4 drops of 1M glycine in Tris buffer at 4°C till immunostaining steps. Before immunostaining, cells were permeabilised in PBS (137 mM NaCl, 2.7 mM KCl, 10 mM Na_2_KPO_4_, 2 mM KH_2_PO_4_, pH 7.3) + 0.1% (v/v) Triton X-100, and subsequently blocked in PBS + 10% normal goat serum (NGS; v/v) + 0.05% Triton X-100.

Immunostaining was carried out in an antibody incubation buffer containing 0.05% Triton and 2% NGS in PBS and coverslips were either attached to microscope slides with cut-out windows, creating a small well where 200 µl of antibody solution (Table 2) was added, or inverted on top of 100 µl antibody solution on a parafilm-covered glass slide. Primary antibodies were added to coverslips and incubated overnight at 4 °C. The samples were then washed with fresh PBS for 3 times in 10-20 min steps, prior to secondary antibody application for 1.5 hrs at room temperature. The samples were then washed in fresh PBS 3 times in 10-20 min steps.

**Table 2.** Antibodies and fluorescent probes used for sample labelling.

| Primary Antibodies | Catalogue number | Source | Dilution |
| --- | --- | --- | --- |
| Anti-EEA1 (Mouse) | 610457 | BD Biosciences, CA | 1:200 |
| Anti-Rab5A (Mouse) | 66339-1-Ig | Proteintech, IL | 1:200 |
| Anti-Rab5A (Rabbit) | 11947-1-AP | Proteintech, IL | 1:200 |
| Anti-endofin (Rabbit) | 13118-2-AP | Proteintech, IL | 1:200 |
| Anti-EGFR1 | Mab 108 | Purified from hybridoma clone HB-9764 [48] | 1:200 |

**Table 2. Antibodies and fluorescent probes used for sample labelling**
| Secondary Antibodies | Catalogue number | Source | Dilution |
| --- | --- | --- | --- |
| Alexa Fluor 488 goat anti-rabbit | A11008 | ThermoFisher Scientific | 1:200 |
| Alexa Fluor 488 goat anti-mouse | A11001 | ThermoFisher Scientific | 1:200 |
| Alexa Fluor 594 goat anti-rabbit | A11012 | ThermoFisher Scientific | 1:200 |
| Alexa Fluor 594 goat anti-mouse | A11005 | ThermoFisher Scientific | 1:200 |
| Atto 647 N goat anti-rabbit | 40839-1ML-F | Merck Life Sciences | 1:200 |
| Atto 647 N goat anti-mouse | 50185-1ML-F | Merck Life Sciences | 1:200 |

### Expansion Microscopy

#### Anchoring

Fluorescently labelled samples were prepared for linking into the gel by incubating them at 4°C overnight in 0.1 mg/ml Acryloyl-X (w/v; Catalogue number A20770, Thermo Fisher Scientific) diluted in 1x PBS.

#### 10x ExM gelation

Cells in each well were washed with PBS for 3 times in 20-minute steps. A gel monomer solution consisted of 4:1 molar ratio of dimethylacrylamide and molecular-grade sodium acrylate, dissolved in deionised H_2_O saturated in N_2_ over 1 hr on ice. Potassium persulfate (KPS; Sigma-Aldrich Ltd, MO) was added at 0.4% molar relative to monomer concentration from a 0.036 g/ml stock, made fresh for each experiment, and the solution was bubbled for another 15 mins on ice. 500 µl of the gel monomer solution was mixed rapidly with 2 µl of TEMED and quickly added to the cells. Gelation was allowed in a sealed acrylic chamber comprising two coverslip walls with the coverslip with cells inverted on top of a gel solution in between these cell walls. Gel polymerisation was completed by > 6 hrs before the gels were extracted carefully from the chambers prior to the digestion step.

#### Digestion and expansion

Gels were subjected to digestion in 0.2 mg/mL proteinase K (ProK, New England Biolabs, MA) dissolved in a “digestion buffer” (50 mM Tris pH 8.0, ThermoFisher; 1 mM ethylenediaminetetraacetic acid (EDTA; Sigma); 0.5% Triton X-100; 0.8 M guanidine HCl (Sigma), in deionized H2O) for 10-13 hrs at RT on an orbital shaker. The gels were washed 3x with PBS for 5-10 minutes at room temperature on an orbital shaker. Gels were either transferred directly to petri dishes for expansion stained with a 1 µg/ml solution of DAPI in PBS for 15-20 minutes, followed by 3 x 10-minute washes in PBS at room temperature on an orbital shaker. Gels were transferred to a large glass petri dish where were expanded by gentle washing in fresh deionized H2O for 30-60 minutes, repeated ∼ 4 times, until there was no further change in the gel size.

#### Variation in 4x ExM (proExM) expansion protocol

The gelation solution was prepared as described previously [21]. Cell samples were incubated with the monomer solution (w/v, Sigma-Aldrich) 8.6% sodium acrylate, 2.5% acrylamide, 0.15% N,N’-methylenebisacrylamide, 11.7% NaCl, PBS, 0.1% ammonium persulfate, and 0.1% N,N,N′,N’-tetramethylethylenediamine first for 30 min at 4°C and then for 2 h at 37°C.

#### Microplate-based ExM sample preparation and pre- /post-ExM imaging

The 3D printed square-well microplates and the laser-cut spacers (described previously [26]) were used for experiments performed for pre- and post-ExM analysis. Cells were cultured at the centre of the 20 x 20 mm square wells. Following fixation and immunolabelling, the spacers were sealed onto the well, leaving a 5 x 5 mm central gap within which the gelation and digestion steps were performed. The spacer was removed prior to hydration of the square well, ensuring that the gel expands without any rotation. In 10-fold ExM experiments, the gel was trimmed down to retain a ∼ 2 x 2 mm corner. The x, y, and z stage coordinates of the cells of interest during pre-ExM imaging were used to calculate and re-locate the structure after expansion (see protocol provided by Seehra et al. [26]; Supplementary methods 3.1) .

### Airyscan microscopy

For imaging ExM samples, the gels were placed within acrylic chambers which were custom-made to fit the stages of the microscopes. The chamber itself was typically square (adapted in size and shape to fit 4x or 10x ExM gels) and consisted of a base made of a glass No. 1.5 coverslip (Menzel Gläser, Germany), custom-coated with 0.1% (v/v) poly-L-lysine (Sigma) in order to retain the gels flush on the coverslip.

All images were obtained on a Zeiss LSM980 AiryScan2 (Carl Zeiss, Jena, Germany) with a Zeiss 40x 1.3 NA oil-immersion Plan Apochromat objective. Imaging was performed in Airyscan mode with the gain and laser power adjusted for each sample to accommodate the fluorescence reduction occurring due to the spatial separation of fluorophores during expansion. Fluorophores were excited using Argon 405 nm and 488 nm, DPSS 561 nm and 642 nm laser lines. Image data was acquired using the Zen Blue interface that allowed us to select the emission bands to minimise spectral cross-talk between dyes. In Airyscan mode, emission was recorded onto the GaAsP detector, and the data were subjected to post-acquisition pixel re-assignment analysis with Zen Blue as described previously [20].

### Negative stain transmission electron microscopy of nanocages

Palladium coated coppers grids coated with carbon film were pre-prepared by the electron microscopy facility team. Carbon film-coated grids were glow discharged for 20-25 seconds and incubated with 3-5 µl of various concentrations of protein sample for 5 – 30 seconds. Grids were blotted and washed 2x by dipping in dH2O and blotting. grids were stained with uranyl formate by a brief wash followed by 20 second incubation, then dried before imaging.

Negatively stained sampled were images with TEM and were examined using a Philips CM100 transmission electron microscope or the Tecnai 12 transmission electron microscope. Magnification was set to 23,000 x. Images were captured using a CCDC camera. The focus was defocused slightly to according to the sample to improve image contrast.

### Image analysis

#### Pre- and post-ExM imaging and distortion analysis

The pre- and post-ExM image volumes were registered to each other using an iterative block-matching analysis pipeline implemented via the plugin Fujiyama via ImageJ (see Supplementary Methods section 3.2 for details). A series of aligned 2D slices from XY and XZ planes throughout the image volumes were then subjected to a Farneback optical flow analysis, described in the Supplementary Methods section 3.3, which produced distortion shift vectors for each 2D slice considered. Pairing up the XY and XZ shift coordinates for a given reference point allowed us to re-calculate the 3D distortion vectors (see visualisation shown in Figure 2C-iii & iv, produced with open-source Mayavi). The analysis and visualisation code is provided in the data supplement.

#### Analysis of nanocages in negative stain TEM image data

Nanocage diameter was determined from electron micrographs. An elliptical boundary was manually defined for each nanocage (as illustrated in Figure 2F-inset). The minor and major axis of each ellipse was determined using the “Measure” function in ImageJ. Nanocage width was determined from the average of the minor and major axis measurements.

#### Detection and analysis of nanocages in Airyscan image data

2D Airyscan image data (unexpanded and post-ExM) were analysed with Python Microscopy Environment (PyME; downloaded from python-microscopy.org on the 11/11/2024). See Supplementary Methods section 3.4 for details.

#### Measurement of endosomal widths in 10x ExM image data

Image volumes were imported into ImageJ and the pixel scale was adjusted to account for either the estimated macroscopic or intrinsic EF. In z-stacks, images where endosomes were in focus were selected and the elipse function was used to manually fit the torus-shaped endosomes. The Measure function was then used to read out the major and minor axes of the elipse and the average of these two values was recorded as the width.

#### Chord plots for displaying marker density and co-localisation

A custom R script (included in Data supplement) was developed to summarise the co-localisation measurements in chord a chord diagram (Figure 4K&L). In each plot, the length of the circumference was sub-divided into arcs assigned to a give protein in proportion to the average proportion of the pixels occupied by the marker at a given time point (as displayed in Figure 4E). The chord were plotted such that the percentage noted at the near end of each chord reported the percentage of that marker co-localising with the marker at the far end.

#### Resolution estimation with PSF measurements

To obtain PSF estimates, 3D stacks were taken of 100 nm multi-spectral polystyrene Tetraspeck Fluorescent microspheres (T7279, Thermo Fisher, UK) suspended in acrylamide. The lateral resolution of the LSM 980 Airyscan 2 system was measured to be ∼ 170 nm.

#### Analysis of endosomal features ExM image data

All puncta localisation, nearest neighbour distance and size analyses were performed in Huygens software (SVI, Hilversum). Manders co-localisation percentages [49] were calculated using the coloc2 plugin in FIJI (ImageJ v1.54f) with the BioFormats plugins. Binary masks for the colocalization analysis were constructed by automatic thresholding using the Moments thresholding option in FIJI. All statistical tests on measurements reported in this paper were non-parametric, and were performed in GraphPad Prism. Open-source software Paraview v6.0.0 was used for 3D isosurface visualisation of data.

The volume of endosomes was traced using a radial sampling approach applied to 3D 10x Airyscan image stacks of endofin (detailed in Supplementary methods section 3.5). This leveraged the observation that endofin appeared to closely outline the endosomes on the cytoplasmic side.

## End Matter

## Author Contributions and Notes

IJ, BC, and PW conceptualised the research project, designed the experiments, and provided the primary resources, reagents, and developed the experimental tools. TBS, RSS, TMDS, and NA performed the experiments. TBS, TMDS, MES, and NA produced the experimental materials. IJ, BC, PW, NFR, and DB provided the supervision. TBS, IJ, RSS, and RK performed the primary data analysis and data curation, TBS, IJ, PW, BC, and NFR interpreted the data. TBS, IJ, PW, and BC wrote the paper.

The authors declare no conflict of interest.

## Acknowledgments

The authors acknowledge the Wolfson Light Microscopy Facility for providing with the microscopes and troubleshooting support for all imaging experiments presented here. This research was funded by UK Research & Innovation and Medical Research Council award made to IJ (MR/S03241X/1), a Wellcome Trust Investigator award to PW (212246/Z/18/Z) and PhD scholarships awarded to IJ from The University of Sheffield’s Faculty of Science and the EPSRC Doctoral Training Programme scheme at Sheffield. The authors extend their gratitude to Prof David Baker and Dr Yang Hsia of The University of Washington for sharing *E. coli* plasmids of the decahedral nanocage variants, and Dr Indrajit Lahiri for advice on characterisation of the nanocages with TEM.

## Notes

### Competing Interest Statement

The authors have declared no competing interest.

## References

1. Chung, I., et al., Spatial control of EGF receptor activation by reversible dimerization on living cells. Nature, 2010. 464(7289): p. 783–7 (10.1038/nature08827).

2. Cullen, P.J. and F. Steinberg, To degrade or not to degrade: mechanisms and significance of endocytic recycling. Nat Rev Mol Cell Biol, 2018. 19(11): p. 679–696 (10.1038/s41580-018-0053-7).

3. Norris, A. and B.D. Grant, Endosomal microdomains: Formation and function. Curr Opin Cell Biol, 2020. 65: p. 86–95 (10.1016/j.ceb.2020.02.018).

4. Rink, J., et al., Rab conversion as a mechanism of progression from early to late endosomes. Cell, 2005. 122(5): p. 735–49 (10.1016/j.cell.2005.06.043).

5. Christoforidis, S., et al., Phosphatidylinositol-3-OH kinases are Rab5 effectors. Nat Cell Biol, 1999. 1(4): p. 249–52 (10.1038/12075).

6. Simonsen, A., et al., EEA1 links PI(3)K function to Rab5 regulation of endosome fusion. Nature, 1998. 394(6692): p. 494–8 (10.1038/28879).

7. Raiborg, C., et al., Hrs sorts ubiquitinated proteins into clathrin-coated microdomains of early endosomes. Nat Cell Biol, 2002. 4(5): p. 394–8 (10.1038/ncb791).

8. Wenzel, E.M., et al., Concerted ESCRT and clathrin recruitment waves define the timing and morphology of intraluminal vesicle formation. Nat Commun, 2018. 9(1): p. 2932 (10.1038/s41467-018-05345-8).

9. Shi, W., et al., Endofin acts as a Smad anchor for receptor activation in BMP signaling. J Cell Sci, 2007. 120(Pt 7): p. 1216–24 (10.1242/jcs.03400).

10. Kazan, J.M., et al., Endofin is required for HD-PTP and ESCRT-0 interdependent endosomal sorting of ubiquitinated transmembrane cargoes. iScience, 2021. 24(11): p. 103274 (10.1016/j.isci.2021.103274).

11. Franke, C., et al., Correlative single-molecule localization microscopy and electron tomography reveals endosome nanoscale domains. Traffic, 2019. 20(8): p. 601–617 (10.1111/tra.12671).

12. van der Beek, J., et al., Quantitative correlative microscopy reveals the ultrastructural distribution of endogenous endosomal proteins. J Cell Biol, 2022. 221(1) (10.1083/jcb.202106044).

13. Huang, B., et al., Three-dimensional super-resolution imaging by stochastic optical reconstruction microscopy. Science, 2008. 319(5864): p. 810–3 (10.1126/science.1153529).

14. Bond, C., et al., Heterogeneity of late endosome/lysosomes shown by multiplexed DNA-PAINT imaging. J Cell Biol, 2025. 224(1) (10.1083/jcb.202403116).

15. Chen, F., P.W. Tillberg, and E.S. Boyden, Expansion microscopy. Science, 2015. 347(6221): p. 543–548 (doi:10.1126/science.1260088).

16. Shi, X., et al., Label-retention expansion microscopy. J Cell Biol, 2021. 220(9) (10.1083/jcb.202105067).

17. Bucur, O., et al., Nanoscale imaging of clinical specimens using conventional and rapid-expansion pathology. Nat Protoc, 2020. 15(5): p. 1649–1672 (10.1038/s41596-020-0300-1).

18. Sheard, T.M.D., et al., Three-dimensional visualization of the cardiac ryanodine receptor clusters and the molecular-scale fraying of dyads. Philos Trans R Soc Lond B Biol Sci, 2022. 377(1864): p. 20210316 (10.1098/rstb.2021.0316).

19. Louvel, V., et al., iU-ExM: nanoscopy of organelles and tissues with iterative ultrastructure expansion microscopy. Nature Communications, 2023. 14(1): p. 7893 (10.1038/s41467-023-43582-8).

20. Sheard, T.M.D., et al., Three-Dimensional and Chemical Mapping of Intracellular Signaling Nanodomains in Health and Disease with Enhanced Expansion Microscopy. ACS Nano, 2019. 13(2): p. 2143–2157

21. Tillberg, P.W., et al., Protein-retention expansion microscopy of cells and tissues labeled using standard fluorescent proteins and antibodies. Nat Biotechnol, 2016. 34(9): p. 987–92 (10.1038/nbt.3625).

22. Wen, G., et al., A Universal Labeling Strategy for Nucleic Acids in Expansion Microscopy. J Am Chem Soc, 2021. 143(34): p. 13782–13789 (10.1021/jacs.1c05931).

23. White, B.M., et al., Lipid Expansion Microscopy. J Am Chem Soc, 2022. 144(40): p. 18212–18217 (10.1021/jacs.2c03743).

24. Lee, H., et al., Tetra-gel enables superior accuracy in combined super-resolution imaging and expansion microscopy. Sci Rep, 2021. 11(1): p. 16944 (10.1038/s41598-021-96258-y).

25. Klimas, A., et al., Magnify is a universal molecular anchoring strategy for expansion microscopy. Nature Biotechnology, 2023. 41(6): p. 858–869 (10.1038/s41587-022-01546-1).

26. Seehra, R.S., et al., Geometry-preserving expansion microscopy microplates enable high-fidelity nanoscale distortion mapping. Cell Reports Physical Science, 2023. 4(12) (10.1016/j.xcrp.2023.101719).

27. Truckenbrodt, S., et al., A practical guide to optimization in X10 expansion microscopy. Nat Protoc, 2019. 14(3): p. 832–863 (10.1038/s41596-018-0117-3).

28. Chozinski, T.J., et al., Volumetric, Nanoscale Optical Imaging of Mouse and Human Kidney via Expansion Microscopy. Sci Rep, 2018. 8(1): p. 10396 (10.1038/s41598-018-28694-2).

29. Damstra, H.G.J., et al., GelMap: intrinsic calibration and deformation mapping for expansion microscopy. Nat Methods, 2023. 20(10): p. 1573–1580 (10.1038/s41592-023-02001-y).

30. Truckenbrodt, S., et al., X10 expansion microscopy enables 25-nm resolution on conventional microscopes. EMBO Rep, 2018. 19(9) (10.15252/embr.201845836).

31. Weisshart, K., The Basic Principle of Airyscanning. Zeiss Technology note, 2014

32. Hsia, Y., et al., Corrigendum: Design of a hyperstable 60-subunit protein icosahedron. Nature, 2016. 540(7631): p. 150 (10.1038/nature20108).

33. Cezanne, A., et al., A non-linear system patterns Rab5 GTPase on the membrane. Elife, 2020. 9 (10.7554/eLife.54434).

34. Horiuchi, H., et al., A novel Rab5 GDP/GTP exchange factor complexed to Rabaptin-5 links nucleotide exchange to effector recruitment and function. Cell, 1997. 90(6): p. 1149–59 (10.1016/s0092-8674(00)80380-3).

35. Stenmark, H., et al., Rabaptin-5 is a direct effector of the small GTPase Rab5 in endocytic membrane fusion. Cell, 1995. 83(3): p. 423–32 (10.1016/0092-8674(95)90120-5).

36. Doyotte, A., et al., The Bro1-related protein HD-PTP/PTPN23 is required for endosomal cargo sorting and multivesicular body morphogenesis. Proc Natl Acad Sci U S A, 2008. 105(17): p. 6308–13 (10.1073/pnas.0707601105).

37. Ali, N., et al., Recruitment of UBPY and ESCRT Exchange Drive HD-PTP-Dependent Sorting of EGFR to the MVB. Current Biology, 2013. 23(6): p. 453–461 (10.1016/j.cub.2013.02.033).

38. Valdes, P.A., et al., Improved immunostaining of nanostructures and cells in human brain specimens through expansion-mediated protein decrowding. Sci Transl Med, 2024. 16(732): p. eabo0049 (10.1126/scitranslmed.abo0049).

39. Damstra, H.G.J., et al., Correction: Visualizing cellular and tissue ultrastructure using Ten-fold Robust Expansion Microscopy (TREx). Elife, 2022. 11 (10.7554/eLife.85169).

40. Tanaka, T., et al., Ligand-activated epidermal growth factor receptor (EGFR) signaling governs endocytic trafficking of unliganded receptor monomers by non-canonical phosphorylation. J Biol Chem, 2018. 293(7): p. 2288–2301 (10.1074/jbc.M117.811299).

41. Rink, J., et al., Rab Conversion as a Mechanism of Progression from Early to Late Endosomes. Cell, 2005. 122(5): p. 735–749 (10.1016/j.cell.2005.06.043).

42. Podinovskaia, M., et al., A novel live-cell imaging assay reveals regulation of endosome maturation. eLife, 2021. 10: p. e70982 (10.7554/eLife.70982).

43. Del Conte-Zerial, P., et al., Membrane identity and GTPase cascades regulated by toggle and cut - out switches. Molecular Systems Biology, 2008. 4(1): p. 206 (10.1038/msb.2008.45).

44. Thevathasan, J.V., et al., Nuclear pores as versatile reference standards for quantitative superresolution microscopy. Nature Methods, 2019. 16(10): p. 1045–1053 (10.1038/s41592-019-0574-9).

45. Votteler, J., et al., Designed proteins induce the formation of nanocage-containing extracellular vesicles. Nature, 2016. 540(7632): p. 292–295 (10.1038/nature20607).

46. Vanheusden, M., et al., Fluorescence Photobleaching as an Intrinsic Tool to Quantify the 3D Expansion Factor of Biological Samples in Expansion Microscopy. ACS Omega, 2020. 5(12): p. 6792–6799 (10.1021/acsomega.0c00118).

47. Eilts, J., et al., Resolving protein organization in cells with nanometer resolution. bioRxiv, 2025: p. 2025.08.11.669713 (10.1101/2025.08.11.669713).

48. Flores-Rodriguez, N., et al., ESCRT-0 marks an APPL1-independent transit route for EGFR between the cell surface and the EEA1-positive early endosome. J Cell Sci, 2015. 128(4): p. 755–67 (10.1242/jcs.161786).

49. Manders, E.M.M., F.J. Verbeek, and J.A. Aten, Measurement of co-localization of objects in dual-colour confocal images. J Microsc, 1993. 169(3): p. 375–382 (10.1111/j.1365-2818.1993.tb03313.x).

