## Supplementary information for "Mapping EGFR1 sorting domains in endosomes with a calibrated 3D expansion microscopy toolkit"

### 1. Supplementary Figures and movies

#### Supplementary Figures

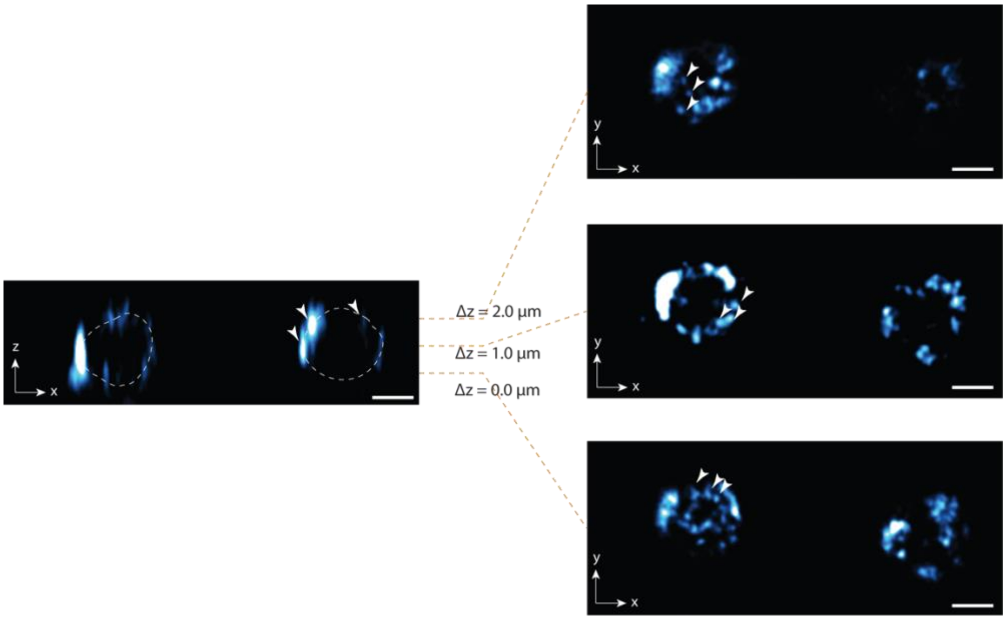


**Supplementary Figure S1: Orthogonal z-x (left) and x-y (right) views of an Airyscan volume of a RPE-1 cell in a region containing two adjacent endosomes.** Shown, are the x-y planes of the top, middle, and bottom planes of the endosomal vesicles in the expanded gels, imaged with an inverted Zeiss LSM 980 Airyscan microscope with a 1.3 NA objective lens. Arrowheads indicate nanoclusters observed on both the lateral, top, and bottom surfaces of the endosomes. Scale bar: 100 nm in pre-ExM length scale.

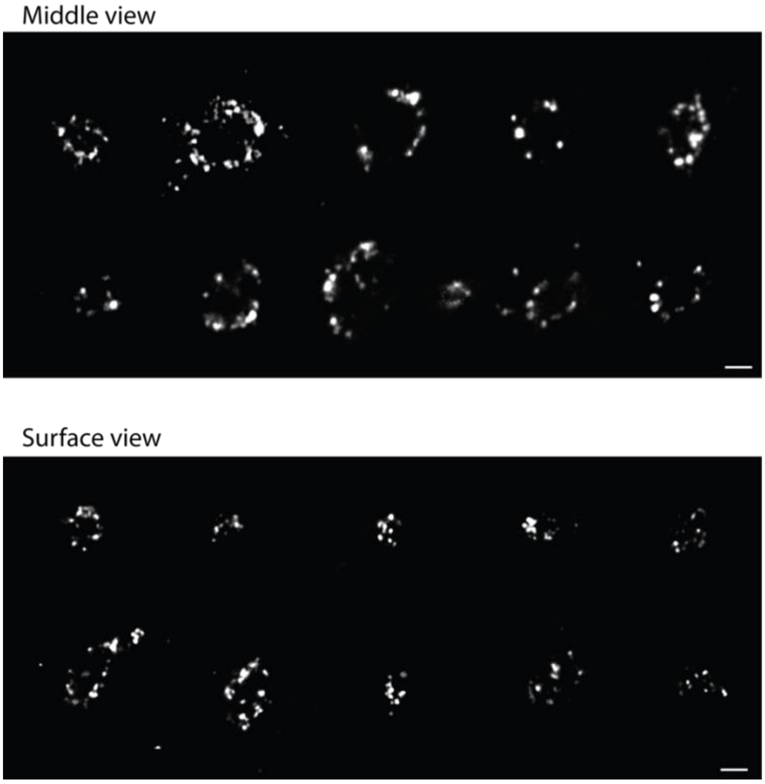


**Supplementary Figure S2: Montage of endosomes in the in-focus middle (top) and glancing surface views (bottom).** Scale bars: 100 nm in pre-expansion length scale.

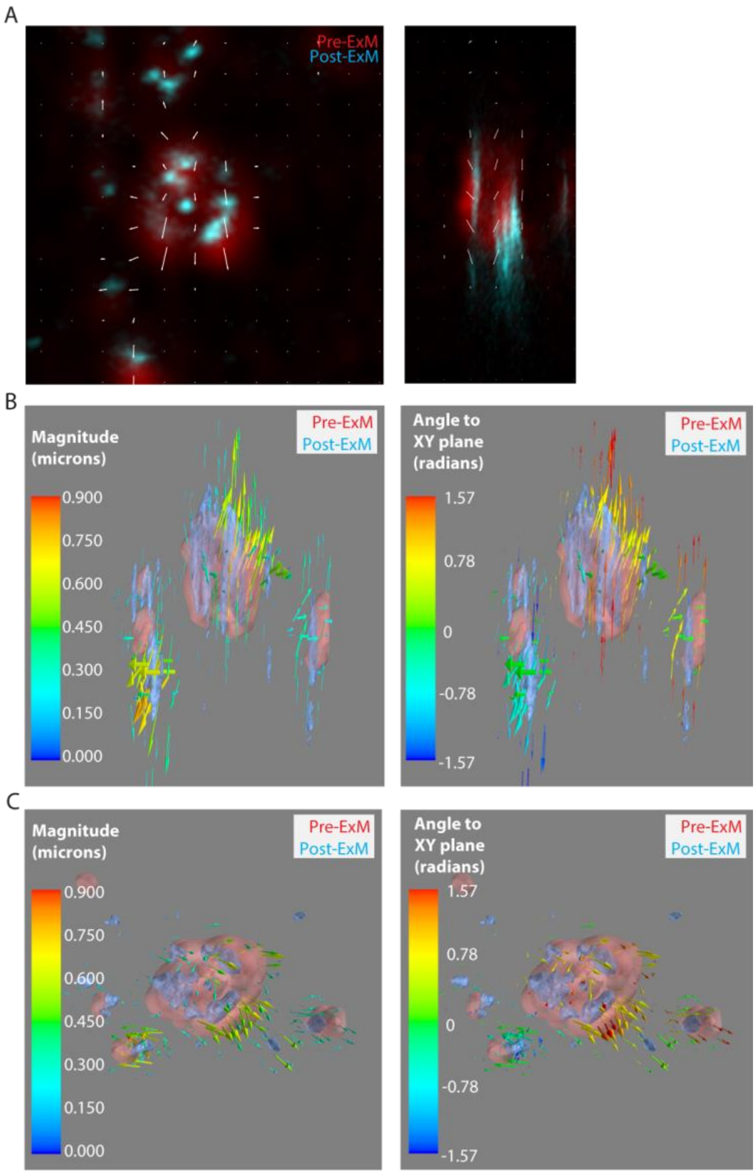


**Supplementary Figure S3: 2D distortion map with comparative 3D viewpoint of an endosome within an EEA-1 fluorophore stained RPE-1 cell.** A) 2D XY and XZ distortion map slices from the centre of the Z-stack. B) 3D XY full volume distortion maps illustrating the uncalibrated magnitude of distortion and the angle of the distortions relative to the XY plane. C) 3D XZ full volume distortion maps illustrating the uncalibrated magnitude of distortion and the angle of the distortions relative to the XY plane.

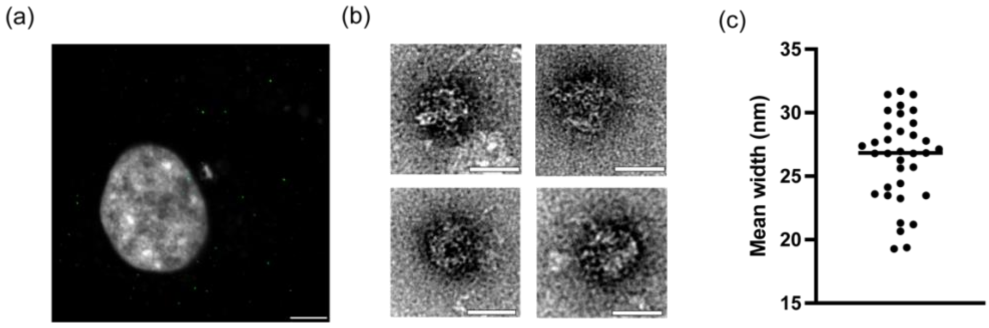


**Supplementary figure S4. Expression and expansion of nanocages in mammalian cells. (a)** Airyscan confocal image of HeLa cell expressing diffraction-limited, fluorescently labelled nanocages (green); scale bar = 5 µm. **(b)** Magnified views of individual nanocages cropped from negative stained transmission electron micrographs of purified lysate of transfected HeLa cells; scale bars = 25 nm. (c) Dot plot of the mean widths measured from the electron micrographs; mean ± SD of 26.4 ± 3.4 nm.

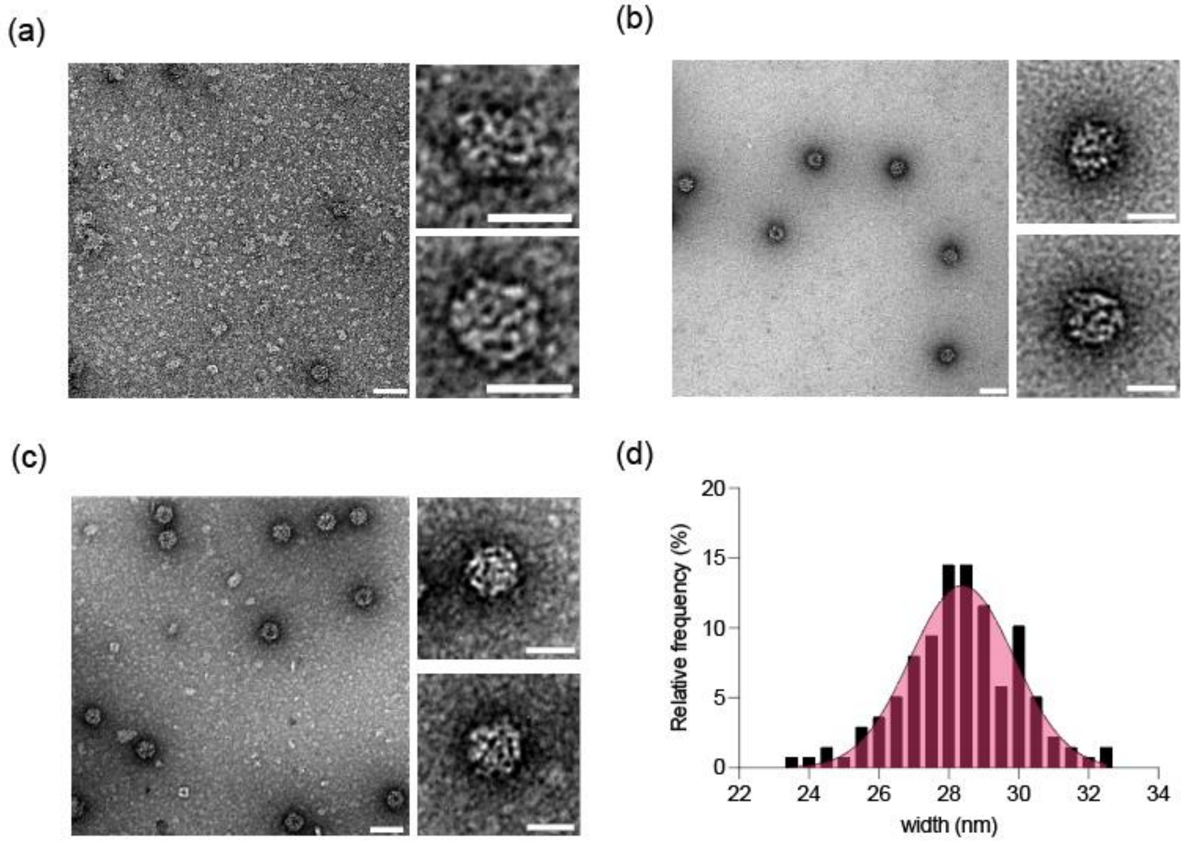


**Supplementary figure S5. Characterisation of nanocages purified from bacterial lysate**. (a) Electron micrograph of nanocages taken directly from IMAC purification. (b and c) Electron micrographs of nanocages after further filtration through a 1MDa membrane via dialysis or spin column concentration. Insets show magnified view of purified nanocages from each dataset. (d) width measurements of nanocages in electron micrographs of post-dialysis samples of the nanocates; n = 138 nm. Scale bars 100nm for wide view and 50 nm for insets.

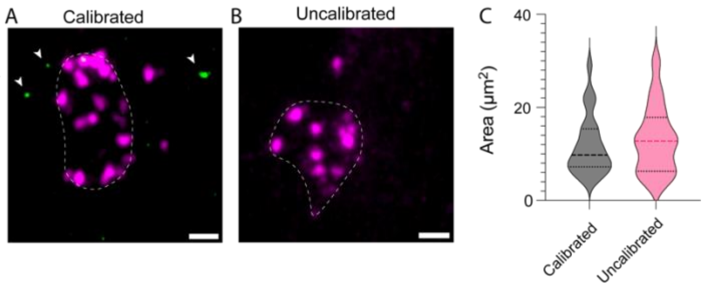


**Supplementary Figure S6: Calibrating intrinsic expansion factor using nanocages.** Maximum intensity projection of 3D images showing EEA1-labelled (A) endosomes (Magenta) alongside calibrating nanocages (green) (left). (B) and an example of a cell that was not transfected with the reporter nanocages (right). White arrows show calibrating nanocages. White dashed lines indicate the EEA1-label boundary. (C) Plots of endosome area measured from the EEA1 boundary in calibrated gels (left, grey) or uncalibrated gels (right, pink). n = 25 endosomes from 8 cells and 4 expansion gels for calibrated measurements; n = 11 endosomes from 9 cells and 4 expansion gels for uncalibrated measurements. Scale bars 100 nm in pre-ExM scale.

### Legends for supplementary movies

**Supplementary movie 1: Surface rendered view of endosomal localisation of EGF and EGFR1:** Exemplar 10x ExM Airyscan image volumes at 15 and 30 min, surface rendered with ParaView 6.0.0. Traced boundary of the endosome shown in translucent grey. EGF and EGFR1 labelling projected onto the traced endosomal surface are shown in cyan blue and magenta respectively.

**Supplementary movie 2: Surface rendered view of endosomal localisation of Rab5a and endofin:** Top and bottom views of 10x ExM Airyscan image volumes surface rendered with ParaView 6.0.0. Rab5a antibody labelling at 30 min is shown in red whilst endofin antibody labelling is shown in yellow. Traced volume of the endosome shown in translucent grey.

### 2. Supplementary Tables

| 15 min | | Protein 2 | | | |
|---|---|---|---|---|---|
| | | Endofin | EGF | EGFR1 | Rab5a |
| Protein 1 | Endofin | | 47.4 ± 18.1% (9) | 69.7 ± 4.6% (5) | 74.5 ± 6.2% (4) |
| | EGF | 65.1 ± 8.8% (9) | | 80.8 ± 8.9% (5) | 57.3 ± 3.9% (4) |
| | EGFR1 | 65.0 ± 8.8% (5) | 81.3 ± 4.8% (4) | | |
| | Rab5a | 70.9 ± 8.7% (4) | 40.3 ± 12.0% (4) | | |

| 30 min | | Protein 2 | | | |
|---|---|---|---|---|---|
| | | Endofin | EGF | EGFR1 | Rab5a |
| Protein 1 | Endofin | | 38.2 ± 16.6% (9) | 61.3 ± 5.8% (4) | 75.0 ± 10.3 % (4) |
| | EGF | 54.2 ± 20.9% (9) | | 75.1 ± 23.5% (4) | 50.6 ± 25.2% (5) |
| | EGFR1 | 54.6 ± 6.9% (4) | 60.0 ± 23.4% (4) | | |
| | Rab5a | 66.0 ± 15.5% (5) | 32.2 ± 13.4 (5)% | | |

**Supplementary Table S1. Summary of co-localisation percentage values at 15 and 30 min time points.** Stated are Mean ± SD (n = number of cells analysed)

### 3. Supplementary Methods

#### 3.1. Plate-based expansion for pre- and post-ExM imaging

##### Expansion microscopy gel preparation and expansion

The geometry-preserving ExM microplates developed previously [1] contained square wells, each with a laser-cut silicone spacer. The gels were polymerised within the inner well created by the spacer. Following gelation, the frame was released and the gel expansion carried out such that the orientation of the gel was preserved. For 4x ExM, the original size of the gel was 5 x 5 mm and it expanded to approximately fit the full well (dimensions: 20 x 20 mm). This workflow is summarised in Supplementary Figure S7. For 10-fold expansion experiments, the hydrogel was trimmed to retain only 2x2 mm at the top left corner prior to expansion.

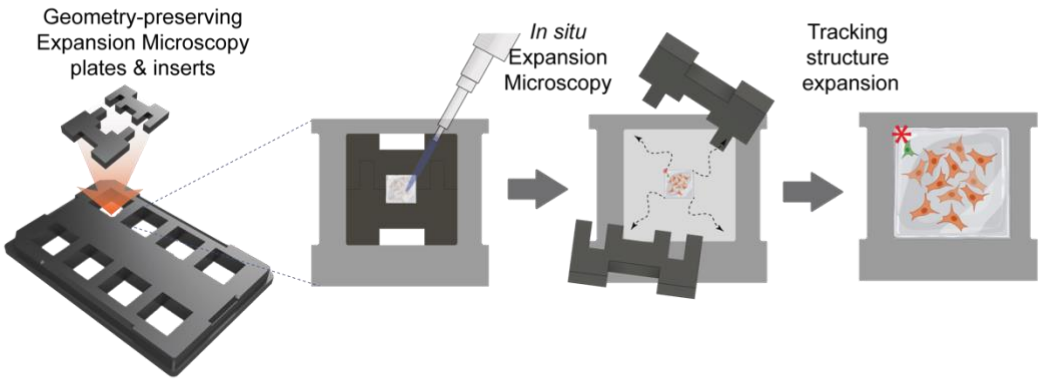


**Supplementary Figure S7. The workflow of the geometry-preserving plate based ExM.**

##### 10-fold ExM recipe adapted for plate wells

Anchoring: Fluorescently labeled samples were anchored to the gel matrix by incubating them overnight at 4°C in 250 µL of PBS containing 0.1 mg/mL of AcX (Acryloyl-X, SE; Thermo Fisher Scientific, A20770), ensuring covalent linkage of amine-containing biomolecules to the polymer network.

Gelation: Cells were rinsed three times with PBS for 20 minutes each to remove residual fixative or media. A gel monomer solution was prepared by combining 1.1 M sodium acrylate (Sigma), 2.0 M acrylamide (Sigma), and 90 ppm (w/v) N,N′-methylenebisacrylamide (bis; Sigma) in deionized $H_2O$. The solution was cooled on ice and deoxygenated by bubbling with nitrogen for 30–60 minutes. Just prior to polymerization, 0.2% (w/v) freshly prepared ammonium persulfate (APS) and 0.2% (v/v) TEMED were added to initiate free-radical polymerization. Approximately 500 µL of gel solution was added to each well, and gelation was performed in a humidified chamber at 37°C for 1–2 hours. After polymerization, the gels were carefully removed for digestion.

Digestion & expansion: Polymerized gels were incubated in digestion buffer (50 mM Tris-HCl pH 8.0, 1 mM EDTA, 0.5% Triton X-100, and 0.8 M guanidine-HCl in deionized $H_2O$) containing 8 units/mL proteinase K (New England Biolabs) for 12–16 hours at room temperature. Following digestion, gels were expanded by repeated washes in excess deionized water, typically three to four times until isotropic expansion stabilized.

##### Coordinate based pre- and post-expansion region tracking

The expected coordinate of the region of interest at post-ExM imaging could be calculated with the following approach using digital stage read-out on *A* (x,y coordinates of the top left corner of the gel during pre-ExM imaging), *B* (x,y coordinates of the region of interest), and *C* (x,y coordinates of the top left corner of the microplate well). Additionally, $f$ (expected expansion factor; i.e. 4) was considered (See Supplementary Figure S8). The pre-ExM vector from A to B ($\overrightarrow{AB}$) was established using the x and y coordinates of the digital stage read-out:

$$\overrightarrow{AB} = B\binom{x}{y} - A\binom{x}{y}$$

Therefore the coordinate of the region of interest (*B′*) during post-ExM imaging was:

$$\overrightarrow{CB'} = \overrightarrow{AB} \times f$$

For example, for a region at pre-ExM of $A = \binom{20}{15}, B = \binom{24}{12}$, and default $f = 4$: $\overrightarrow{AB} = \binom{4}{-3}$.

Therefore $\overrightarrow{CB'} = \binom{4}{-3} \times 4 = \binom{16}{-12}$.

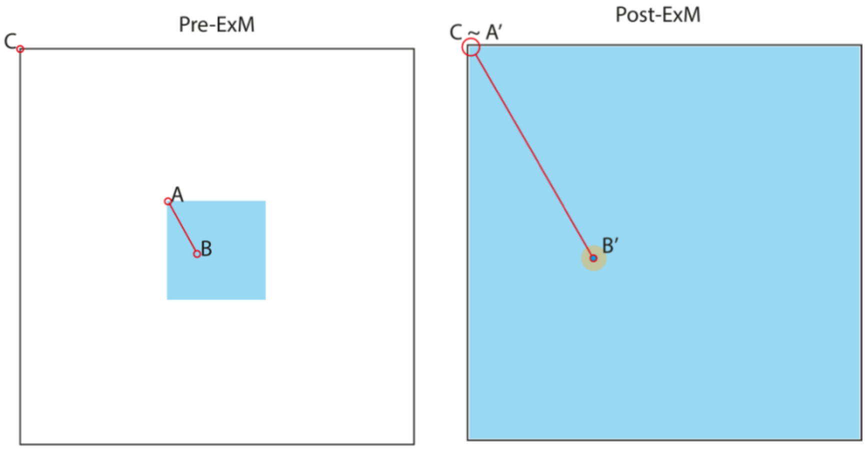


**Supplementary Figure S8:** Principle of Coordinate-based tracking of region of interest during pre-ExM and post-ExM imaging. At pre-ExM imaging, the x and y coordinates of locations A (top left corner of gel), B (location of the region of interest), and C (top left corner of well) are recorded. During post-ExM imaging, the top left corner of the gel (A') is expected to reach the top left corner of the well (C). Calculation of $\overrightarrow{CB'}$ (red line in Post-ExM gel) therefore returns the stage to the x,y coordinates of the expanded region of interest. Figure adapted from Seehra et al. (2023).

#### 3.2. Pre- and post- ExM 3D image alignment

Image data volumes were initially drift-corrected as required using ImageJ's "Linear Stack Alignment with SIFT" [2] and pre-ExM data was adjusted by multiplying the pixel size by an approximation of the expansion factor. This was performed to simplify the initial alignment process.

Both datasets were opened with the Fijiyama [3] plugin in ImageJ, "Two images registration (training mode)" approach. The plugin allows for this volumetric assessment and employs the block-matching algorithm [4] for 3D volume alignment. This process uses 'blocks' composed of voxels of the structure to find matching structural information to aid in the alignment. The resulting output was then assessed to calculate a global transformation that best explains the local correspondences (Supplementary figure S9).

To facilitate the process an initial manual alignment of the data was performed, with preference given to structures closer to the coverslip. This was done to reduce the influence of any optical aberrations that may be present in deeper z planes of the sample. The block-matching algorithm used the similarity-based tools: rotation, translation, and uniform scaling. Hence, the resulting transformation matrix contains data reflective of the 3 axes present. However, consideration for how well the images are aligned was determined by accuracy to 1-2% of the overall image features, a factor dependent on the initial alignment.

Initial alignments were conducted manually, using the similarity and 2D viewer settings. Here, landmarks were selected and matched between pre- and post-ExM image volumes. Image data was primarily aligned based on structures closest to the coverslip to reduce bias towards optically induced effects that may occur higher into the dataset.

To address any bias here, a standardised approach was adopted. Firstly, the data would be initially aligned within Fijiyama [3] with preference to structures near the coverslip. Secondly, the automatic block-matching algorithm with similarity was employed using the default settings. Finally, the aligned volumes were viewed to determine if the alignment had occurred and whether it needed to be repeated. This final step was a quality control step to ensure that the data showed alignment in the expected orientation. If automatic alignment resulted in a significant misalignment from the manually aligned data, such that few similar features remained in close proximity, the process was repeated with greater care to align the initial data (Supplementary figure S10).

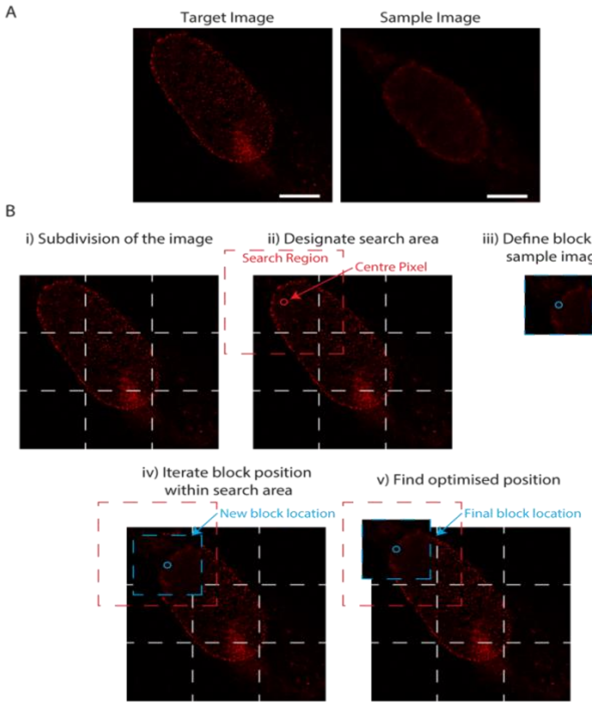


**Supplementary Figure S9: Block-matching process for two-image matching, a 2D representation of the 3D process.** Example illustrated with pre- and post-ExM volumes of a nucleus with Nups98 staining. A) Assignment of the target (post-ExM) and sample (pre-ExM) images. B) i) The target and sample images are subdivided into blocks (voxels). ii) A central pixel and a corresponding search volume are designated in the target image. iii) The equivalent block from the sample image is defined. iv) The sample block is iterated within the 3D search volume. v) Shift and scaling information regarding the optimised position is recorded. This process is repeated for all blocks.

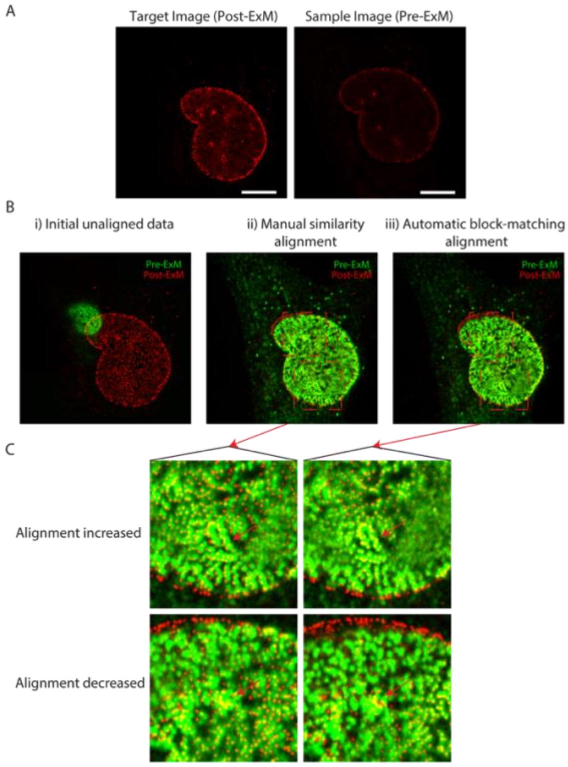


**Supplementary figure S10: Example workflow of Fijiyama usage.** A) Designating the pre-/post ExM images. B) i) Initial presentation of unaligned pre-ExM (green) and post-ExM (red) data. ii) Initial manual registration using similarity alignment. iii) Aligned data after automatic block-matching algorithm with similarity. C) Representative images showing alignment changes between manual and automatic approaches.

#### 3.3. 3D distortion analysis

##### Validation of Farneback Optical Flow 2D Approach in 3D ExM

As per 2D distortion analysis, Farneback's optical flow algorithm [5] was used to assess distortions. To incorporate 3D data, the algorithm was applied to the comparative 2D XY and XZ planes of the image sets to compile the distortion data. Comparisons of the distortion data of the shared axis from XY and XZ were used to validate the process. The absolute sum of the shared X axis distortion data was the same between the two planes.

##### Application of 3D Distortion Mapping with Comparative 2D Regions

Using this approach, the distortion vectors were used alongside the pre-/post-ExM image data to produce visualisations of the distortions present. This 3D visualisation is a representation of the 3D surface render of the data. By limiting the vector amount visualised the distortions were observable.

The observability of distortions arises from both the difference in pre- and post-ExM overlays, and the magnitude of the local distortion vectors. The distortion vectors were colour coded and scaled by the magnitude of the vectors (Supplementary Figure S11).

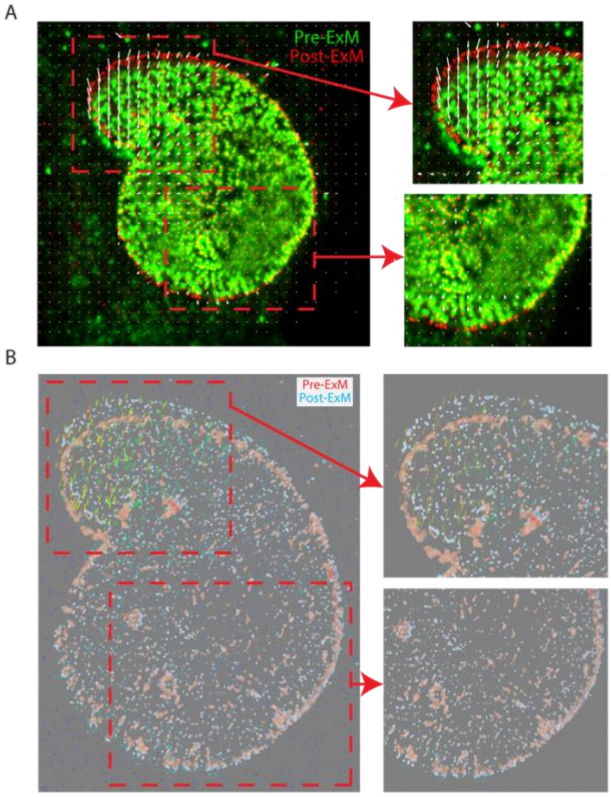


**Supplementary figure S11: 2D distortion map with comparative 3D viewpoint of a RPE-1 cell nucleus stained with NUP98.** A) 2D distortion map with sections illustrating high and low distortion. B) 3D distortion map in the same plane wherein the size and colour intensity of the distortion vectors illustrate high and low regions of distortion.

##### RMSE as a Function of Length Measurement Calculations for 3D Distortion Data

To quantify the impact of 3D distortion vector information on the overall error in the post-ExM images, 3D RMSE analysis code (RMS code 3D_V3.py) was developed. In particular, the 2D RMSE code developed previously [1] used a list of coordinates as its input. These were subsequently processed using linear algebra to acquire the distance measurements required to calculate the RMSE. This list of coordinates contained only X and Y values.

The numpy.linalg.norm [6] function performed at order = 2 was equivalent to the Euclidean distance. Therefore, the 3D implementation of the analysis was achieved by amending the list of coordinates to contain the coordinates for all three dimensions.

#### 3.4. Detection and analysis of nanocages in Airyscan image data

3D Airyscan images with in-focus nanocages were selected for the analysis. The images were imported into dh5view data viewer in Python Microscopy Environment, accessible freely from python-microscopy.org. On the viewer, the modules, 'blobFinding' and 'blobMeasure' were selected from the Modules dropdown menu. The following settings were defined on the 'Object finding' panel before clicking the 'Find' command. Once the spots were detected, The 'Object fitting' panel was used to perform a 3D Gaussian adaptive fitting command by clicking 'Fit'. Table S2 below summarises the parameters were used for pre- and post-ExM images.

| Parameter | Pre-ExM | Post-ExM |
|---|---|---|
| Object Finding | | |
| Threshold | 10.0 (subjective to an appropriate background intensity level) | 6.0 (up to 10.0) |
| Threshold mode | Multi-threshold | Multi-threshold |
| Channel | Channel 0 | Channel 0 |
| Blur size | 4.0 | 6.0 |
| Object Fitting | | |
| ROI half-size | 8 | 12 |

**Supplementary Table S3. Analysis parameters used for detecting and fitting the nanocage spots using Python Microscopy Environment.** Stated are Mean ± SD (n = number of cells analysed)

The output of Object Fitting consists of the variable, 'wxy' which reflects the in-plane sigma.

To export coordinates and fitting values to a .txt file, the command Save > Save fit results was used. To deduce the full-width at half-maximum (FWHM) of the nanocages fitted, the sigma values were multiplied by 2.3548.

#### 3.5. Endosome volume reconstruction

##### 3.5.1. Segmentation and 3D reconstruction

To reconstruct the full volume of endosomes from labels that localised to its limiting membrane, we developed a method that segments, smooths and 3D renders an approximated surface. This method relies on a single image channel consisting of a full 3D z-stack of a target that localises to the limiting membrane (e.g. endofin shown in supplementary figure S12-A).

The 10x Airyscan z-stack of endofin labelling was sub-sampled to capture the z range that spans the height of the endosome. It was important to crop the image volume to contain the labelling on the endosome only due to the influence that background or non-specific labelling densities bear on the boundary estimation of the endosome. In the example shown, regions of background or non-endosomal labelling (see inset of supplementary figure S12-A) was cropped out. Once, cropped the volume was zero-padded on each size by 20 voxels and analysed with custom-written Python script, Delaunay_3D_volume.py (included in the data supplement; see section 3.5.3. for protocol).

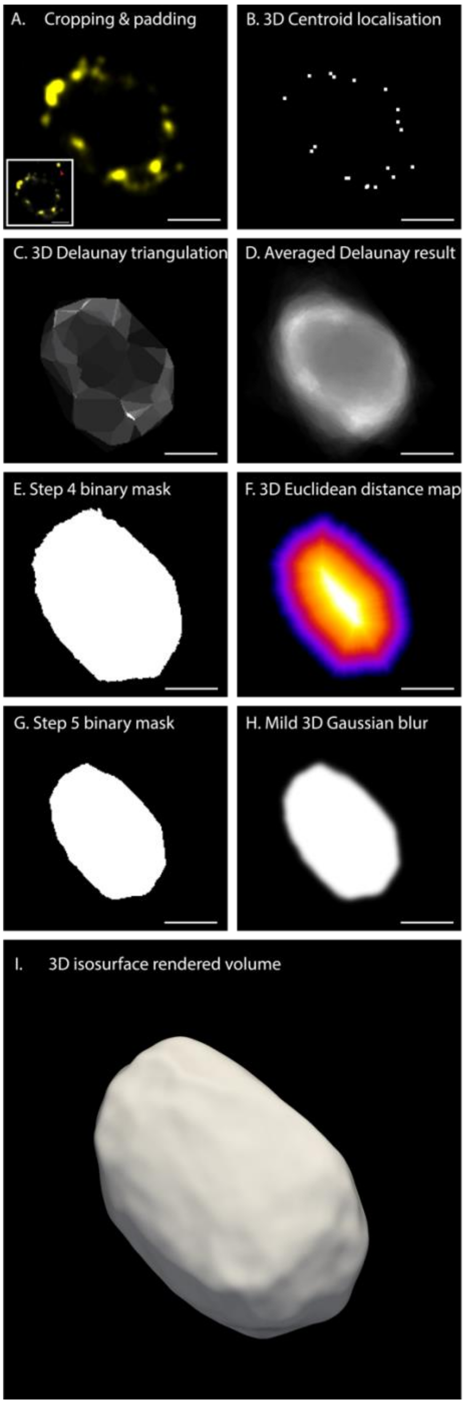


**Supplementary figure S12: Endosomal surface tracing with endofin labelling** A) 10x ExM Airyscan z stack was selected due to the localisation of endofin near the surface of the endosome. Background labelling were cropped out and the image volume was zero-padded by 20px on each side. Example shows each step in a single 2D x-y slice through the middle of the stack. B). Result of the centroid detection of the puncta of endofin labelling, shown as a shallow projection across a depth of 100 nm (corrected for EF). C) 3D Delaunay triangulation result where each triangle was given an intensity value inversely proportional to its longest side. D). The resulting image from averaging 40 Delaunay triangulation calculations where, in each iteration, the coordinates were jittered by a distance between 0 and 20 x fitting error. E) binary mask constructed by linear intensity thresholding of the image in panel D such that the labelling puncta are aligned with the endosome boundary. F) Euclidean distance transform constructed from the mask in panel E. G) The final binary mask of the endosome volume, calculated by thresholding the distance transform. H) A smoothed endosome volume achieved by convolving the volume in panel G by a 3D Gaussian with a sigma ~ 0.7 px in each dimension. I) 3D isosurface rendered image of the smoothed endosome volume in panel H. Scale bar: 200 nm (corrected for expansion factor)

Briefly, the image volume was subjected to an automated difference of Gaussian (DOG) background suppression step. The punctate labelling densities were then subjected to localisation analysis that constituted the steps (i) detection of puncta, (ii) fitting with a 3D Gaussian model, (iii) ablation of the detected punctum from the volume to prevents re-detection. Supplementary figure S12-B shows a shallow projection of the centroids of the endofin puncta of the shown example. The output of the fitting analysis included the x, y, and z coordinates as well as a fitting error value (in nm) for each punctum. The localised coordinates were then used to construct a 3D Delaunay triangulation in a series of between 40 and 300 independent iterations. Prior to the Delaunay triangulation calculation in each iteration, each coordinated was jittered by a 3D displacement determined by a random uniform probability between 0 and a maximum proportional to the fitting error. Each triangle resulting from the Delaunay triangulation was assigned a fill value linearly proportional to the longest side of that triangle (supplementary figure S12-C). The iterations of the Delaunay triangulation were then averaged (supplementary figure S12-D) and an intensity threshold is applied (supplementary figure S12-D). In the thresholding step, the user is prompted to select the lowest threshold that allows the convex shape of the endosome to be captured whilst including the puncta *within* that volume. Depending on the overall density and coverage of the labelling across the surface of the endosome, resultant mask (supplementary figure S12-E) provides a good approximation of the surface topology and the shape of the endosome. However, the segmentation strategy that dictates such that labelling is included *within* the segmented object, the overall volume of the resulting mask was often larger than the endosome. A second thresholding step was carried out in order to preserve the shape and topology of the endosome whilst matching the volume to the localisation of the puncta. For this step, a 3D Euclidean distance map (EDM) was constructed within the segmented volume (supplementary figure S12-F). The user was then prompted to select a threshold that allows the EDM to be eroded down to a volume such that the puncta line the new surface of the endosome (supplementary figure S12-G).

For the smooth isosurface rendering of the resultant mask, it was convolved by a small Gaussian PSF (with a sigma of ~ 0.7 pixels in each dimension; supplementary figure S12-H). The smoothed mask was then imported to ParaView v6.0.0. and isosurface rendered (supplementary figure S12-I).

##### 3.5.2. Repeated volume tracing with independent endosomal markers

The reproducibility of the volume traced with this method relies upon the accuracy of and the density of the label chosen for tracing the endosome. To examine variability to introduced to the volume reconstruction from different markers, we considered two types of doubly labelled 10x ExM datasets.

In the first example (supplementary figure S13-A), we compared BODIPY630 NHS ester staining and anti-EEA1 labelling patterns of the same endosome. Being a nondescript stain of lipid-rich compartments [7], BODIPY630 staining returned a dense and volumous image of the endosome. By comparison, EEA1 reported distinct puncta that predominantly localised to the outline of the endosome. The coverage of the EEA1 labelling across the endosome was considerably lower, whilst some puncta also appeared to extend beyond the endosome and needed to be cropped out of the volume tracing analysis. Being a nondescript stain, BODIPY630 also produced a weak, but nonzero background stain in the surrounding neighbourhood which required exclusion by cropping. 91-99% of the voxels of the reconstructed volumes were in agreement (supplementary figure S13-A-iv, v, vi).

By comparison, the volume reconstructions from two independent labels, EGF-Alexa488 and anti-EGFR1 (supplementary figure S13-B), that reported similar densities and punctate morphologies of labelling showed much stronger agreement (~ 92-100% overlap).

###### *Limitaions to consider prior to use*

The reliance on Delaunay triangulation allow the interpolation of the endosomal boundary between unlabelled regions. This feature makes it unsuitable for tracing endosomes which feature either tubulation or invagination. Another inherent assumption of the method is that the chosen marker of the endosomal boundary is indeed present at the boundary. Whilst the use of a reference label (e.g. membrane reporter or an NHS ester stain) is advisable, a minimum expectation is that the chosen marker is retained at high density along the limiting membrane of the endosome. Endofin was selected as the marker of choice in the analyses presented in Figure 4, however it was only possible because the density of endofin was sufficiently high at the two time points and the culture conditions considered.

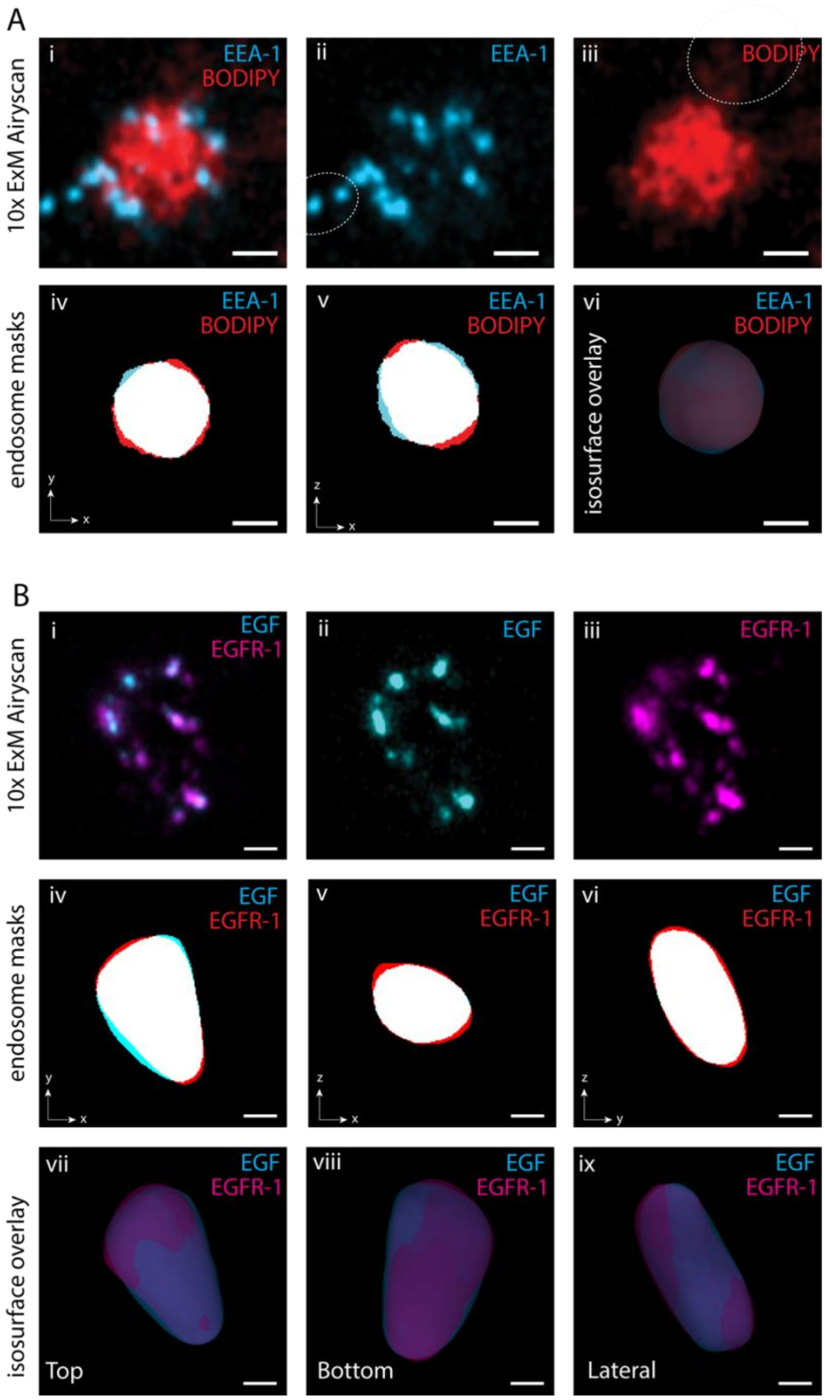


**Supplementary figure S13: Independent volume tracing with multiple markers**. (A-i) An example of a 10x ExM Airyscan image volume of an endosome stained for EEA1 (cyan) and BODIPY630 NHS ester (red). (A-ii, iii) Single channel images of EEA1 and BODIPY respectively. Dashed lines indicate surrounding labelling cropped out manually prior to the volume tracing analysis. (A-iv, v) Overlays of the independently binarised endosomal volumes from EEA1 (cyan) and BODIPY (red) in x-y and x-z views. 97.9% of EEA1 and 92.1% BODIPY voxels were found to overlap with the opposite volume model. (A-vi) Surface rendered overlay of the two independent volumes traced with EEA1 (cyan, translucent), and BODIPY (red, translucent). (B-i-iii) Overlay and individual channels of a 10x ExM Airyscan volume of an endosome stained for EGF (cyan) and EGFR1 (magenta). (B-iv, v, vi) Overlays of the independently binarised endosomal volumes from EGF (cyan) and EGFR1 (red) in x-y, x-z, and y-z views. 99.9% of EGF and 90.8% EGF voxels were found to overlap with the opposite volume model. (B-vii-ix) Top, bottom, and lateral views of isosurface rendered overlay of the two independent volumes traced with EGF (cyan, translucent), and EGFR1 (red, translucent). Scale bars, 100 nm (in pre-ExM scale).

##### 3.5.3. Mapping endosomal surface labelling onto reconstructed surface

Two methods were used for mapping labelling densities outlining the endosomes in 10x ExM volumes onto the reconstructed surface of the endosomal volume (see supplementary figure S14-A overlaying the reconstructed surface voxels overlaid with the ExM image of endofin). This was achieved by directly projecting the labelling densities onto the surface layer of voxels of the reconstructed endosomal volume. In both approaches, the Normal axis to each voxel of the reconstructed surface was considered.

In the first approach, the maximum voxel value across 40 nm either side of the Normal axis was then assigned to the surface voxel (supplementary figure S14-B). In effect, this produced a maximum intensity projection of the ExM image on the reconstructed endosomal surface (supplementary figure S14-C).

In the second approach, the image volume of the protein of interest was subjected to a centroid detection analysis (equivalent to the puncta fitting analysis in section 3.5.1). The voxel of the reconstructed surface was then marked for the detected centroid and convolved with a 3D Gaussian model with sigma matching the Gaussian fitted in that step. The resultant image produced a series of smooth objects that were overlaid against the isosurface visualisation of the reconstructed volume (supplementary figure S14-D).

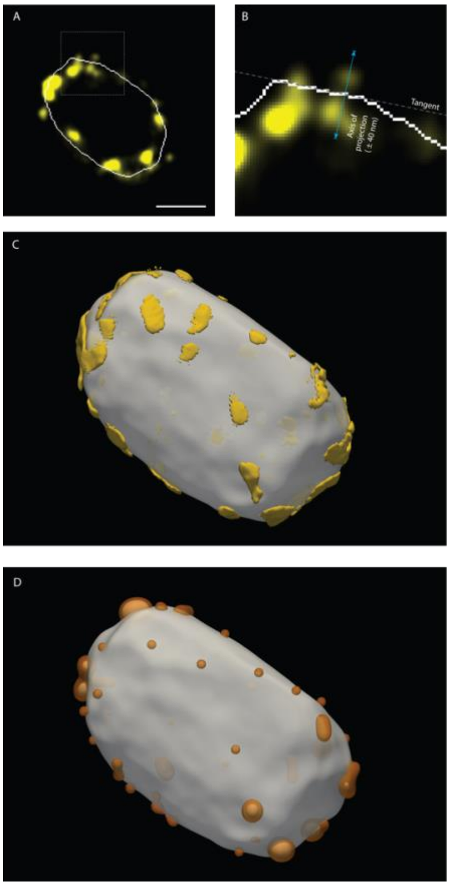


**Supplementary figure S14: Projecting proteins of interest onto traced endosome.** A) In the shown example, an image volume of an endosome labelled with endofin was considered (yellow). Overlaid in white, is the boundary voxel layer of the traced volume. Example shows one 2D x-y slice through the middle of the image volume whilst the analysis wans conducted in 3D. B) On each traced surface voxel, the axis Normal to the local surface region (blue line) was analysed. Of the intensity values of the protein channel (yellow) on 40 nm either side of the surface along this Normal axis, the maximum voxel value as assigned to the considered voxel of the traced surface. C) The volume containing this maximal intensity projection (yellow) was isosurface rendered together with the overall traced volume of the endosome (grey). D) As an alternative approach, punctate densities of the protein of interest were localised using a 3D Gaussian fitting method. In this view, the fitted Gaussian profiles were projected onto the endosome volume surface and isosurface rendered (orange). Scale bar, 200 nm.

##### 3.5.3. Protocol for endosomal volume reconstruction and surface mapping

1. A computer with ~ 32 Gb of RAM and computational performance equivalent to an Apple M4 Macbook Pro or a Windows 11 PC with a Ryzen 7 5700U with speed upto 4.3GHz is a minimum requirement for this analysis.
2. Ensure a Python v3.10 (or later) installation including the most common libraries including numpy, pandas, tifffile, matplotlib, scipy, and skimage.
3. In the working folder, ensure the following files are present:

- Delaunay_3D_volume.py (provided script for volume tracing)
- Project_puncta_to_surface.py (provided script for mapping endosome labelling to surface)
- Stack-1.tif (3D ExM image volume of the endosomal labelling)
- Stack-2.tif (Protein of interest for surface mapping; same as stack-1 if the original protein is being mapped)

4. On the terminal, copy into the working directory for example,

```
cd /Users/z3541596/analysis_directory/
```

5. Execute the python script:

```
Python Delaunay_3D_volume.py
```

6. The following window will be displayed. To select the threshold most appropriate for tracing the endosomes, pick the 'thr' value of the example where the smallest traced volume that encompasses the puncta (in the x-z views shown). In this example, the optimal threshold value is 0.00398. Close the window and enter this value into the command line prompted and press Return.

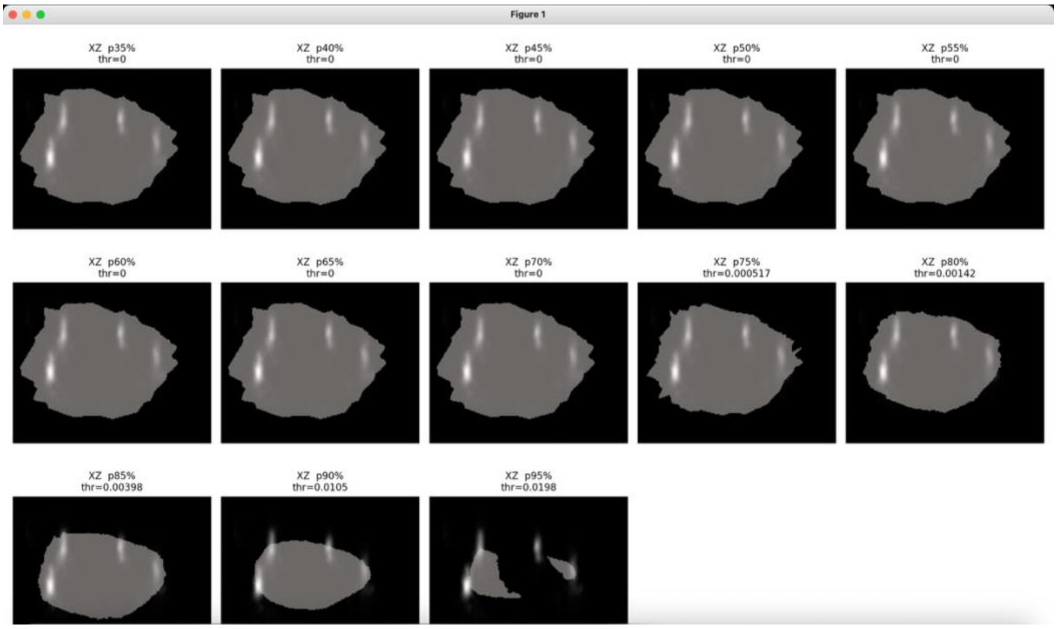


7. You will then be prompted to select an additional threshold in the following window. Pick the threshold that best connects the puncta (i.e. avoid the threshold that encompass the puncta because this over-estimates the size of the endosome). In the example shown, a 'thr' value 17.5 is optimal. Close the window and enter this value into the command line prompted and press Return.

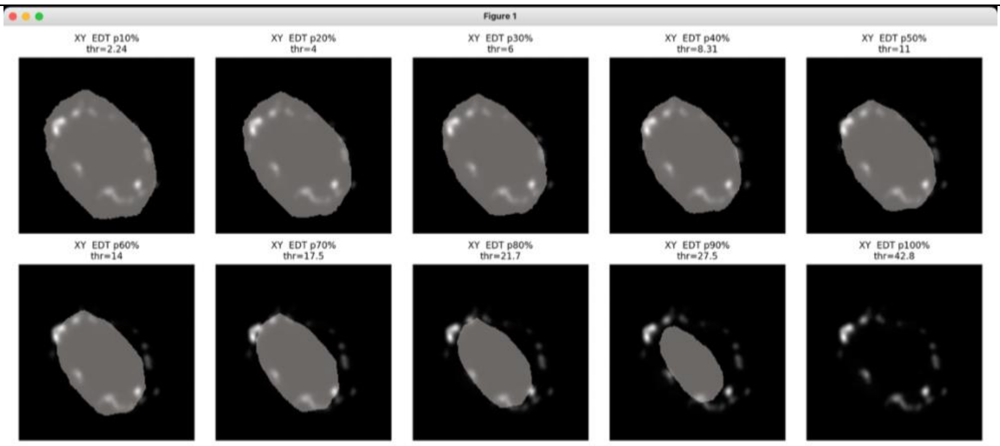


8. Ensure that the following files are present in the working directory:
- Protein.tif (3D ExM image volume of endosomal labelling of interest)
- Stack-2.tif (raw image volume used for volume tracing)
- Step5_distance_mask.tif (traced cell volume mask – final output from step 3)

9. Execute the python script:

```
Project_Puncta_to_Surface.py
```

10. For 3D rendering, open ParaView (freely available to download from https://www.paraview.org/) and open image volumes listed below (screenshot of example included below).

11. step5_distance_mask_blurred.tif Endosomal volume for isosurface reconstruction
12. puncta-max-projected-Gauss.tif Labelling of interest project onto volume surface
13. puncta-discretised-projected [optional] 3D Gaussian rendered puncta following detection

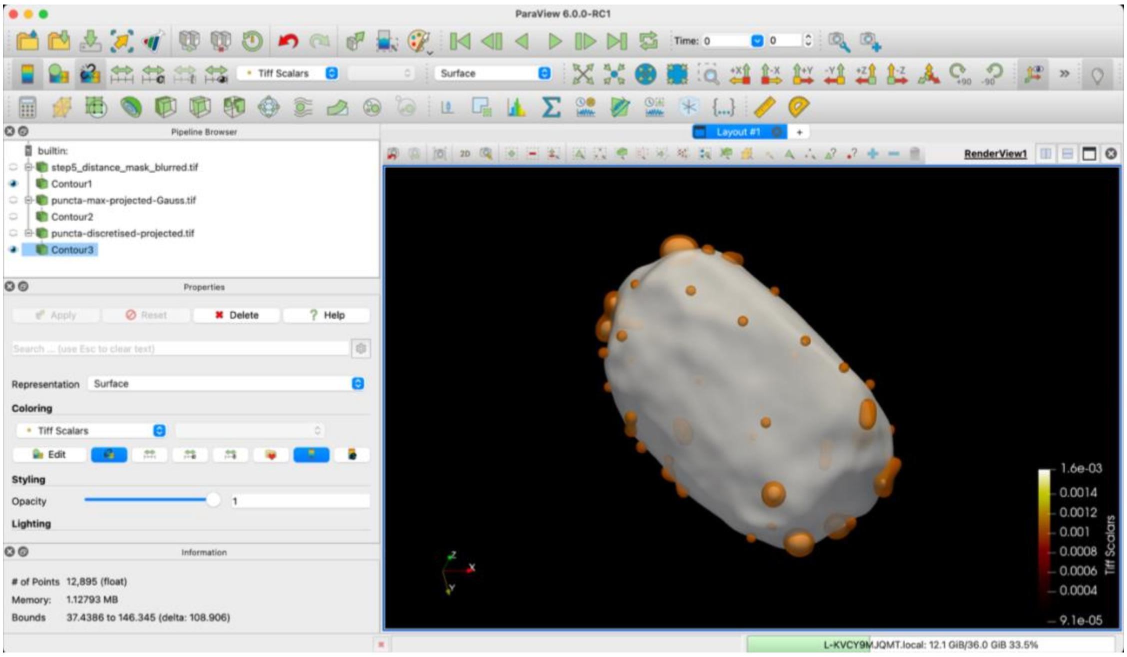

### 4. References


1. Seehra, R.S., et al., *Geometry-preserving expansion microscopy microplates enable high-fidelity nanoscale distortion mapping.* Cell Reports Physical Science, 2023. **4**(12).
2. Lowe, D.G., *Distinctive Image Features from Scale-Invariant Keypoints.* International Journal of Computer Vision, 2004. **60**(2): p. 91-110.
3. Fernandez, R. and C. Moisy, *Fijiyama: a registration tool for 3D multimodal time-lapse imaging.* Bioinformatics, 2021. **37**(10): p. 1482-1484.
4. Hassen, W. and H. Amiri. *Block Matching Algorithms for motion estimation*. in *2013 7th IEEE International Conference on e-Learning in Industrial Electronics (ICELIE)*. 2013.
5. Farnebäck, G. *Two-Frame Motion Estimation Based on Polynomial Expansion*. in *Image Analysis*. 2003. Berlin, Heidelberg: Springer Berlin Heidelberg.
6. Harris, C.R., et al., *Array programming with NumPy.* Nature, 2020. **585**(7825): p. 357-362.
7. Sheard, T.M.D., et al., *Differential labelling of human sub-cellular compartments with fluorescent dye esters and expansion microscopy.* Nanoscale, 2023. **15**(45): p. 18489-18499.
